# PBP1 of *Staphylococcus aureus* has multiple essential functions in cell division

**DOI:** 10.1101/2021.10.07.463504

**Authors:** Katarzyna Wacnik, Vincenzo A Rao, Xinyue Chen, Lucia Lafage, Manuel Pazos, Simon Booth, Waldemar Vollmer, Jamie K Hobbs, Richard J Lewis, Simon J Foster

**Affiliations:** School of Biosciences, University of Sheffield, Sheffield, UK; The Florey Institute for Host-Pathogen Interactions, University of Sheffield, Sheffield, UK; Biosciences Institute, Newcastle University, Newcastle upon Tyne, UK.; Department of Physics and Astronomy, University of Sheffield, Sheffield, UK; Centre for Bacterial Cell Biology, Bioscience Institute, Newcastle University, Newcastle upon Tyne, UK.

**Author notes:** To whom correspondence should be sent. The Royal Society for the Protection of Birds, The Lodge, Sandy, Bedfordshire SG19 2DL (Richard J Lewis).

## Abstract

Bacterial cell division is a complex process requiring the coordination of multiple components, to allow the appropriate spatial and temporal control of septum formation and cell scission. Peptidoglycan (PG) is the major structural component of the septum, and our recent studies in the human pathogen *Staphylococcus aureus* have revealed a complex, multi-stage PG architecture that develops during septation. Penicillin binding proteins (PBPs) are essential for the final steps of PG biosynthesis – their transpeptidase activity links together the peptide sidechain of nascent glycan strands together. PBP1 is required for cell division in *S. aureus* and here we demonstrate that it has multiple essential functions associated with its enzymatic activity and as a regulator of division. Loss of PBP1, or just its C-terminal PASTA domains, results in cessation of division at the point of septal plate formation. The PASTA domains can bind PG and thus coordinate the cell division process. The transpeptidase activity of PBP1 is also essential but its loss leads to a strikingly different phenotype of thickened and aberrant septa, which is phenocopied by the morphological effects of adding the PBP1-specific *β*-lactam, meropenem. Together these results lead to a model for septal PG synthesis where PBP1 enzyme activity is responsible for the characteristic architecture of the septum and PBP1 protein molecules coordinate cell division allowing septal plate formation.

## Introduction

Peptidoglycan (PG) is the major structural component of the bacterial cell wall and is essential for maintaining cell shape, integrity and survival (Silhavy et al., 2010; Turner et al., 2014; Vollmer et al., 2008). The final assembly stages of assembly of this large polymeric molecule are mediated by penicillin-binding proteins (PBPs), key PG synthases that, through their transglycosylase (TG) and transpeptidase (TP) activities, polymerise glycan chains and cross-link them into a mesh-like hydrogel (Pasquina-Lemonche et al., 2020; Typas et al., 2011). Since the cell wall is essential for maintaining bacterial life, PBPs and PG synthesis are a target of some of the most important antibiotics, *β*-lactams (penicillins) and glycopeptides (vancomycin) (Schneider & Sahl, 2010; Zapun et al., 2008). The major human pathogen *Staphylococcus aureus* has a minimalist PBP system as it encodes only four PBPs, PBP1 to PBP4 (Pinho et al., 2013). Only PBP1 (class B PBP with only TP activity, bPBP) and PBP2 (class A bifunctional PBP with both TG and TP activities, aPBP) are essential and sufficient for septal and peripheral PG synthesis in *S. aureus* (Lund et al., 2018; Pinho et al., 2013). PBP2 is the major PG synthase of *S. aureus*, and the septal formation activity of PBP2 is mediated by its substrate, Lipid II (Pinho & Errington, 2005). Although PBP2 is essential, loss of its TP activity can be compensated for by a horizontally acquired class B PBP2A in methicillin-resistant *S. aureus* (MRSA) (Pinho, Filipe, et al., 2001). PBP2A, however, cannot replace PBP1, whose loss is detrimental to the viability of *S. aureus* (Pereira et al., 2007). PBP1 and PBP3 form cognate pairs with the monofunctional TGs, FtsW and RodA, belonging to the SEDS (shape, elongation, division and sporulation) family (Meeske et al., 2016) to facilitate septum formation (PBP1-FtsW) and to maintain the prolate cell shape (PBP3-RodA) of *S*. *aureus*, respectively (Reichmann et al., 2019). PBP4 is a class C PBP with D,D-carboxypeptidase activity (cPBP) and has a TP activity that contributes to the high-level cross-linking of PG and MRSA resistance to *β*-lactams (Loskill et al., 2014; Srisuknimit et al., 2017).

Although *S. aureus* PBPs have been studied over many years, the specific roles of PBP1 in cell division, PG synthesis and architecture have remained elusive. Previous studies have shown that whilst PBP1 is essential, its TP activity is not, implying another role (Pereira et al., 2007; Reichmann et al., 2019). However, this work was performed in an MRSA background that contains PBP2A, encoded by *mecA*, which is non-native to *S. aureus* (Pinho, de Lencastre, et al., 2001). Whilst PBP2A cannot replace PBP1, how these proteins interact is unknown. We have recently shown that the presence of *mecA* has a profound effect on cellular physiology (Panchal et al., 2020). Thus it is important to understand individual and combined roles of *S. aureus* PBPs both in the presence and absence of the exogenous PBP2A, as the vast majority of *S. aureus* infections are caused by methicillin sensitive strains.

## Results

### *S. aureus* PBP1 PASTA domains are essential for growth and PBP1 functionality

PBP1 has a cytoplasmic N-terminal region, a membrane spanning sequence, an exocytoplasmic dimerization domain and a C-terminal region consisting of the TP domain and two PASTA domains (for <u>p</u>enicillin-binding protein <u>a</u>nd <u>s</u>erine/threonine kinase <u>a</u>ssociated domain) (Yeats et al., 2002). We created a set of conditional mutants of *pbp1* to investigate the role of PBP1 in cell division and PG synthesis. An ectopic copy of *pbp1* under the control of the P*spac* promoter (P*_spac_*-*pbp1*) was placed at the lipase locus (*geh*::P*_spac_*-*pbp1*) of *S. aureus* SH1000, and a series of changes were made in this genetic background at the native *pbp1* locus: (i) an in-frame deletion of *pbp1* (*Δpbp1)*, (ii) a deletion of the region encoding the two PASTA domains (*pbp1Δ*_PASTA_), and (iii) the substitution of the catalytic Ser314 to Ala in the TP domain (*pbp1\**) (Fig. 1a, b). We examined the essentiality of PBP1, the PASTA domains and the active TP domain with these mutants. Depletion of PBP1 via IPTG removal (Fig. 1c and Fig. 1 – figure supplement 1a, b) resulted in cell death, confirming the essentiality of PBP1 (Fig. 1c, d and Fig. 1 – figure supplement 1c, d,).

**Fig. 1.**
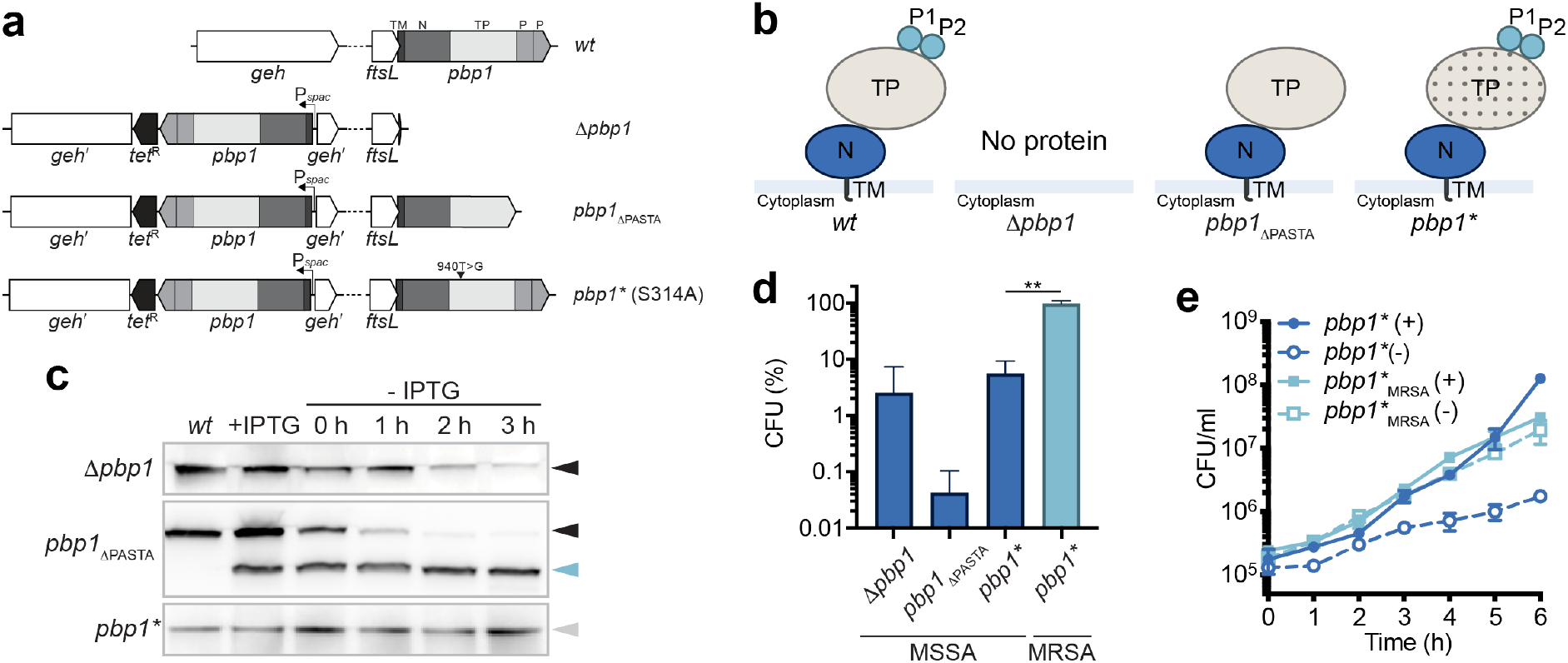
Essentiality of PBP1. **a**, Schematic representation of genetic constructs used in this study. In *S. aureus* WT (*wt*) the 5′ end of *pbp1* overlaps with the 3′ of *ftsL*. The *pbp1* gene encodes a protein containing the transmembrane helix (TM), N-terminal domain (N), transpeptidase (TP) domain and two PASTA domains (P1 and P2). In the mutants, an ectopic copy of *pbp1* is placed under the control of the P*spac* promoter at the lipase (*geh*) locus, whereas the gene in the native *pbp1* locus is either deleted (*Δpbp1*), has P1 and P2 domains removed (*pbp1_Δ_*_PASTA_), or has a point mutation which results in inactivation of the TP domain (*pbp1\**). **b**, Schematic representation of domain architecture of PBP1 in *S. aureus* WT (*wt*) and PBP1 forms produced by *Δpbp1*, *pbp1_Δ_*_PASTA_ and *pbp1\** mutants in the absence of inducer. The TP domain inactivation is shown by dotted shading. **c**, Immunoblot showing PBP1 levels in SH1000 *lacI* (*wt*) and in *Δpbp1*, *pbp1_Δ_*_PASTA_ and *pbp1\** grown with IPTG (+IPTG) and for 0, 1, 2 and 3 h without inducer (-IPTG) analysed using anti-PBP1 antibody. Expected sizes: PBP1 and PBP1* = 83 kDa (black and grey arrowheads, respectively) and PBP1_ΔPASTA_ = 67 kDa (light blue arrowhead). **d**, Plating efficiency of *Δpbp1*, *pbp1_Δ_*_PASTA_ and *pbp1\** (MSSA) cells and MRSA *pbp1*\* (*pbp1*\*_MRSA_) cells upon inducer removal compared to the control groups grow in the presence of inducer. *P* value was determined by Mann–Whitney *U* tests. *P* = 0.0043 (**, *P* < 0.01). Data represent mean ± SD. **e**, Growth curves of *pbp1*\* (MSSA) and MRSA *pbp1*\*(*pbp1*\*_MRSA_) in the presence (+) or absence (-) of IPTG. Data represent mean ± SD. Error bars that are smaller than the symbols are not shown. Data are representative of three (c, e) and at least four (d) independent experiments.

Deletion of the PASTA domains also led to growth inhibition and more than 99% cell death within 4 h (Fig. 1d and Fig. 1 – figure supplement 1c, d). Importantly, this phenotype was not associated with PBP1*_Δ_*_PASTA_ instability (Fig. 1c and Fig. 1 – figure supplement 1a) or loss of its ability to bind its substrate analogue BocillinFL (Fig. 1 – figure supplement 1b). By contrast, deletion of the PASTA domains of *Streptococcus pneumoniae* PBP2x, a PBP1 orthologue, resulted in a complete loss of BocillinFL binding (Maurer et al., 2012). These results indicate the PASTA domains are essential for *S. aureus* growth and PBP1 functionality but not its enzymatic activity.

During construction of the *pbp1\** mutant we obtained, by serendipity, a *pbp1*_STOP_ mutant in which a SNP in the codon for Glu292 resulted in its replacement with a premature stop codon and the truncation of the entire TP and PASTAs region of PBP1 (Fig. 1 – figure supplement 1e, f). However, immunoblot analysis using anti-PBP1 sera could not confirm the presence of the PBP1_STOP_ protein in the *pbp1*_STOP_ mutant (Fig. 1 – figure supplement 1g), suggesting that stability of the N-terminal domain of PBP1 is dependent on its C-terminus.

Although inactivation of PBP1 TP activity (PBP1*) did not affect protein stability (Fig. 1c), it did remove the ability of PBP1 to bind BocillinFL (Fig. 1 – figure supplement 1b). The loss of PBP1 TP activity resulted in severely compromised growth on solid media (Fig. 1d and Fig. 1 – figure supplement 1c) and reduced cellular viability in liquid culture (Fig. 1e and Fig. 1 – figure supplement 1d). Thus, the TP activity of PBP1 is required for growth in the SH1000 background. Inactivation of the PBP1 TP activity was reported previously not to affect growth in the COL strain background (Reichmann et al., 2019). The differences in the necessity for the PBP1 TP activity could result from COL being MRSA whereas SH1000 is a methicillin sensitive *S. aureus* (MSSA).

### PBP1 TP activity is crucial in MSSA but not in MRSA

We have recently developed a set of defined strains where high-level *β*-lactam resistance of MRSA is mediated by *mecA* encoding PBP2A and a mutation in either *rpoB* or *rpoC* (Panchal et al., 2020). This combination of genetic alterations (*mecA^+^ rpoB*) are present in COL (Panchal et al., 2020). To test if the apparent disparity in PBP1 role is associated with MRSA, we developed a high-level resistant mutant of *pbp1\** in the well-characterised *S. aureus* SH1000 by adding the *mecA rpoB*^H929Q^ to the MSSA *pbp1\** mutant, resulting in SH1000_MRSA_ *pbp1\** (Fig. 1 – figure supplement 2a). Inactivating PBP1 TP did not affect the ability of SH1000_MRSA_ *pbp1\** to grow in the absence of IPTG, whereas *pbp1* depletion led to growth inhibition in the isogenic *Δpbp1* MSSA and MRSA strains (Fig. 1d, e and Fig. 1 – figure supplement 2b-d). Thus, the fundamental role of PBP1 in growth and division can only be studied in an MSSA background as otherwise the role of PBP1 can be confounded by the presence of the MRSA resistance apparatus.

### PBP1 PASTA domains are required for septum progression

PG synthesis still occurred in *Δpbp1*, *pbp1_Δ_*_PASTA_ and *pbp1\** in the absence of IPTG, despite cell growth inhibition, as measured by the incorporation of the fluorescent D-amino acid derivative HADA (Fig. 2a). This was not a consequence of the non-synthesis, exchange reaction carried out by PBP4 as it occurred in *pbp4* as well as with the dipeptide ADA-DA (Kuru et al., 2019; Lund et al., 2018) (Fig. 2 – figure supplement 1). All variants increased in cell volume upon depletion of *pbp1*, whereas *pbp1_Δ_*_PASTA_ was enlarged by almost twice as much as *Δpbp1* and *pbp1\** (Fig. 2a, b and Fig. 2 – figure supplement 2a). Despite differences in cell size, both *Δpbp1* and *pbp1_Δ_*_PASTA_ decreased the number of cells with complete septa (Fig. 2a, c). Transmission electron microscopy (TEM) showed that more than 80% of the population had growth defects including cell wall thickening, PG blebs, mis-shapen and/or multiple, incomplete septa. (Fig 2d, e and Fig. 2 – figure supplement 2b, c). Such septa had abnormally thick bases and sharply pointed leading edges, suggesting that cell growth arrest was not due to a lack of septal initiation but instead arrest of inward septum progression.

**Fig. 2.**
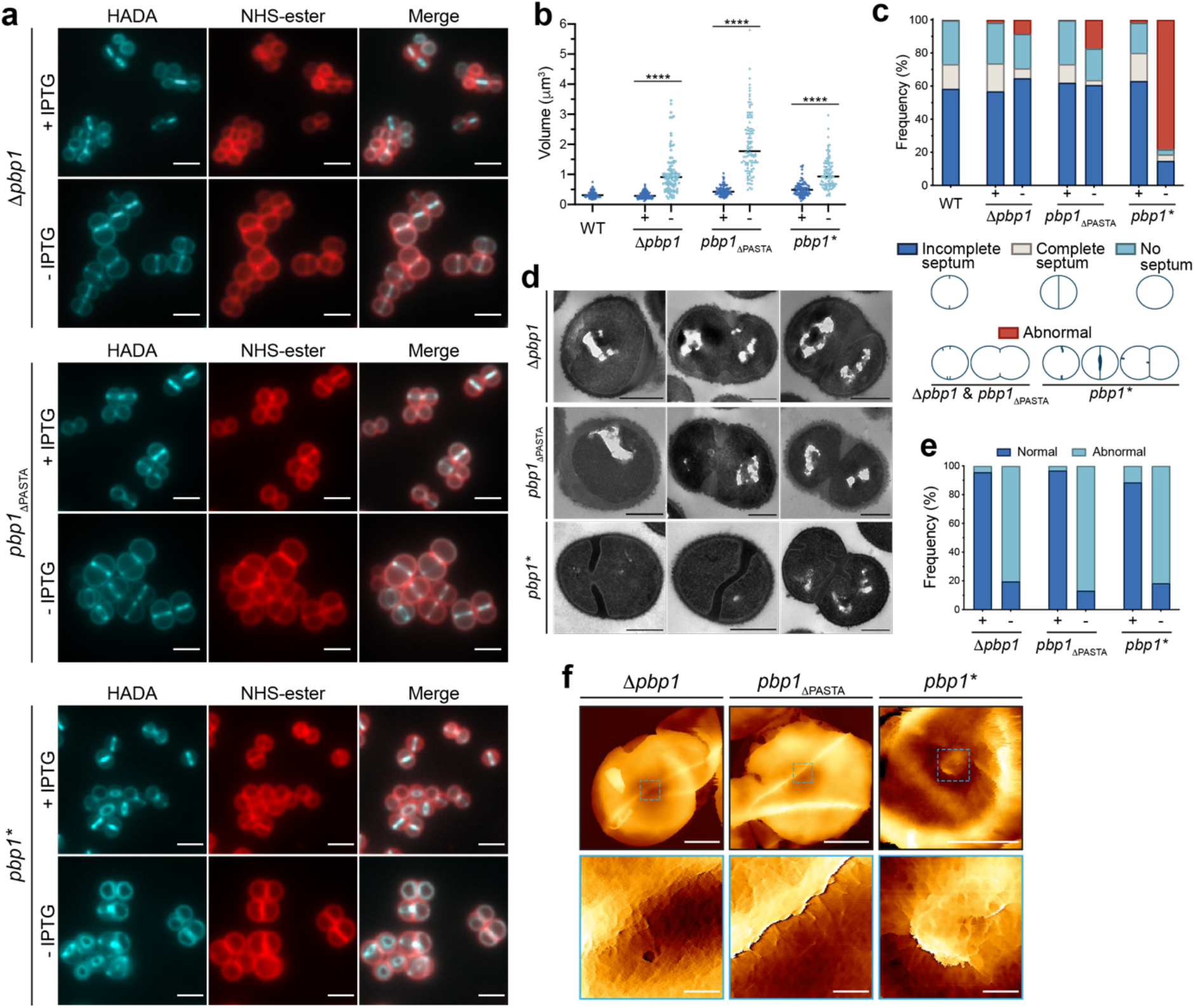
The role of PBP1 in cell division and PG synthesis in *S. aureus*. **a**, Δ*pbp1, pbp1*_ΔPASTA_ and *pbp1\** grown with or without IPTG for 2 h, incubated with HADA for 5 min to show nascent PG, and counter-labelled with NHS-ester Alexa Fluor 555 to image the cell wall. Images are average intensity projections of *z* stacks. Scale bars 2 μm. **b**, Cell volumes of WT (*wt*), Δ*pbp1, pbp1*_ΔPASTA_ and *pbp1\** grown with (+) or without (-) IPTG for 2 h as measured by fluorescence microscopy after NHS-ester Alexa Fluor 555 labelling. Each dot represents a single cell. The median of each distribution is indicated by a black line. The number of cells analysed for each mutant and condition was *n* ≥ 100. *P* value was determined by Mann–Whitney *U* tests (****, *P* < 0.0001). From left to right: *P* = 3.033e- 033, 4.670e-049 and 2.206e-022; *n* = 100, 101, 100, 101, 100, 100 and 101. **c**, Quantification of cellular phenotypes for WT *(wt),* Δ*pbp1, pbp1*_ΔPASTA_ and *pbp1\** based on HADA incorporation (Fig. 2a) after incubation with (+) or without (-) IPTG for 2 h. From left to right *n* = 370, 427, 332, 314, 364, 512 and 331. **d**, TEM of Δ*pbp1, pbp1*_ΔPASTA_ and *pbp1\** grown for 2 h in the absence of inducer. Scale bars 500 nm. **e**, Quantification of cellular phenotypes based on TEM data of Δ*pbp1, pbp1*_ΔPASTA_ and *pbp1\** grown for 2 h in the presence (+) or absence (-) of IPTG. Examples of cells classified as normal (blue) are shown in Fig. 2 – figure supplement 2b. Cells with abnormal phenotypes (light blue) are shown in Fig. 2d and Fig. 2 – figure supplement 2c. From left to right *n* = 391, 329, 314, 377, 263 and 302. **f**, AFM topographic images of internal surface of purified sacculi from Δ*pbp1, pbp1*_ΔPASTA_ and *pbp1\** grown in the absence of inducer for 2 h. Sacculi (top images, scale bars 500 nm, data scales (*z*): 450, 300 and 100 nm from left to right, respectively) and higher magnification images (bottom images, scale bars 50 nm, data scales (*z*): 70, 100 and 50 nm from left to right respectively) scanned within the boxed areas from the top images. Data are representative of two (d-f) and (a-c) three independent experiments.

Atomic force microscopy (AFM) has revealed previously that the first step in cell division is the formation of a PG feature called the “piecrust”, prior to the septal plate(Turner et al., 2010). The *S. aureus* septal plate has two PG architectures: disordered mesh facing the cell membrane and concentric rings in the septum core (Pasquina-Lemonche et al., 2020). Here lack of PBP1 or the PBP1 PASTA domains led to formation of more than one, and often misplaced, piecrust. These mutations also caused an increase in unfinished septal annuli and alterations in the ring surface architecture (Fig 2f and Fig. 2 – figure supplement 3a, c – arrowheads), a characteristic feature of the division plane, freshly revealed immediately after cell scission (Pasquina-Lemonche et al., 2020). Thus, depletion of PBP1 did not stop septum initiation but the loss of the PASTA domains was enough to cause formation of irregular piecrusts, arrest septal plate formation and lead to an altered septal PG architecture.

### PBP1 TP activity regulates septal PG architecture

The *pbp1\** mutant gave a novel phenotype quite distinct from loss of entire PBP1 or the PASTA domains. Inactivation of PBP1 TP activity did not prevent initiation and closing of the septa, but instead resulted in accumulation of cells with aberrant septa and separation defects in about 80% of the population (Fig. 2a, c, e). The septa in such cells had a rounded leading edge, were curved, abnormally thick (Fig. 2d, e and Fig. 2 – figure supplement 2b, c), had agglomerations of mesh-like material close to the septal centre in addition to irregular piecrusts as observed by AFM (Fig. 2f and Fig. 2 – figure supplement 3a, b). The intracellular agglomerations are PG as they stain heavily with HADA and ADA-DA (Fig. 2a and Fig. 2 – figure supplement 1c, f) and could be observed in purified sacculi (Fig. 2f and Fig. 2 – figure supplement 3b). No ring architecture, only mesh structured PG could be observed on the surface of the *pbp1\** mutant. Importantly, using fluorescence microscopy the *pbp1* pbp3 pbp4* mutant, in which PBP2 is the only active TP, presented a similar phenotype upon IPTG removal as *pbp1\**, exemplified by misshapen septa and agglomerations of PG material marked by HADA (Fig. 2 – figure supplement 2d). Therefore, septal synthesis and progression still occurred in the *pbp1\** mutant, however, they resulted from PBP2 transpeptidase activity and potentially the transglycosylase activity of FtsW.

The *pbp1\** phenotype occurred specifically because of the loss of the TP activity of this essential enzyme. This phenotype is mirrored by the mode of action of *β*-lactam antibiotics, which bind to and inhibit the TP activity of PBPs (Schneider & Sahl, 2010). Our results suggest that PBP1 TP activity has a specific role in septal plate formation and without this the septum is mis-shapen. The conditional lethal strains made here allow for functional analysis of the genes concerned. However, phenotypes tend to accumulate on depletion of the wild-type protein over time confusing the precise roles for individual components. To independently corroborate the role of the TP activity of PBP1 we utilized an approach to directly, and selectively, inhibit its activity. Meropenem (MEM) has a higher affinity for PBP1 than PBP2 (Berti et al., 2013; Yang et al., 1995) and, therefore, we hypothesised that its effect on *S. aureus* would match *pbp1\**. In a MEM-titration, treatment with 1x MIC MEM was sufficient to lead to cell death and a significant increase in SH1000 WT cell volume after 1 h (Fig. 3a, b and Fig. 3 – figure supplement 1a). More than 70% of MEM treated cells had growth defects that manifested as aberrantly shaped septa and accumulation of PG as shown by HADA labelling (Fig. 3a, c, d Fig. 3 – figure supplement 1c, e), similar to observations made with the *pbp1\** mutants (Fig. 2a, c-e, Fig. 2 – figure supplement 1c, f). The MEM phenotype of malformed septa was not linked to PBP3 or PBP4 as it was also observed in the corresponding double mutant (Fig. 3c, d and Fig. 3 – figure supplement 1b, d, f), which corroborated the role of PBP2 in misshapen septal genesis.

**Fig. 3.**
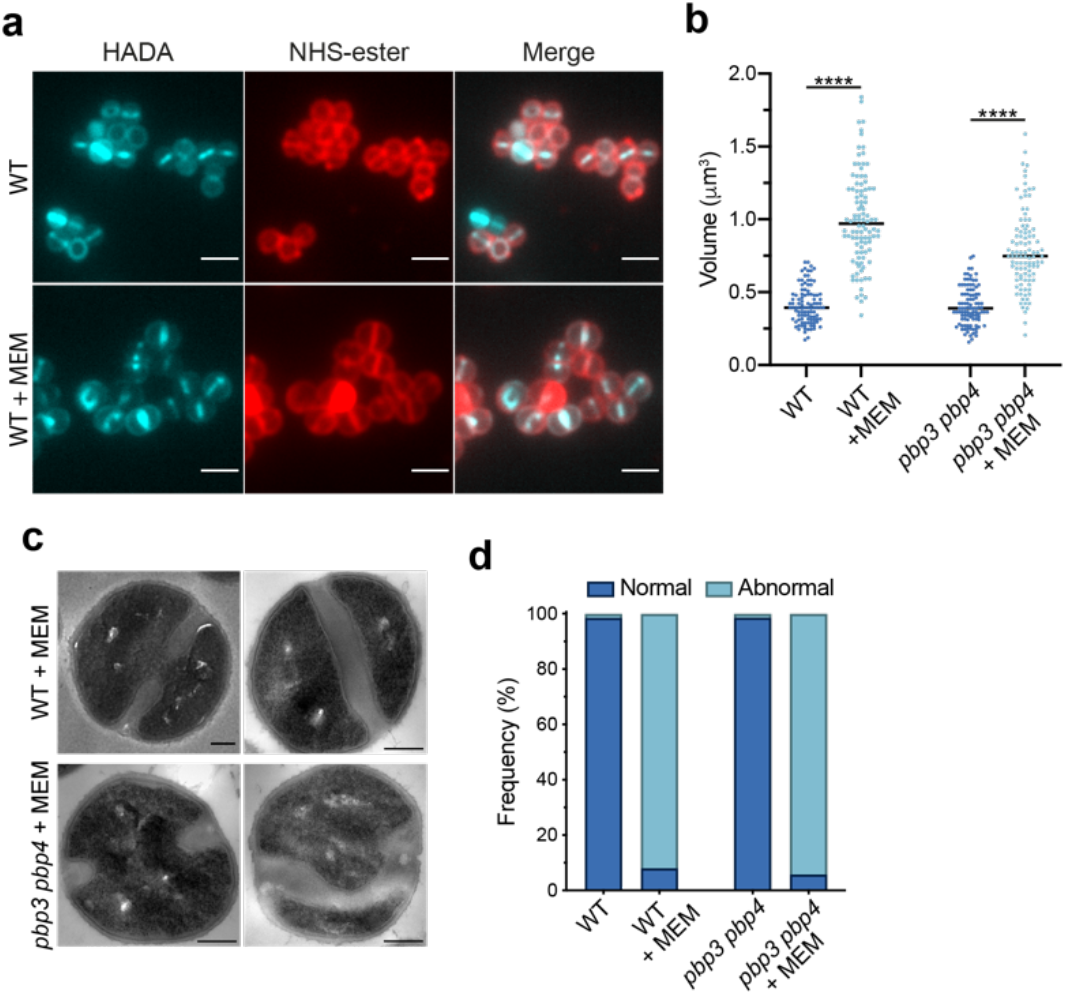
Effect of meropenem (MEM), an antibiotic with high affinity for PBP1, on *S. aureus*. **a**, Fluorescence images of SH1000 WT treated with 1x MIC MEM for 1 h, labelled with HADA for 5 min to show nascent PG and counter labelled with NHS-ester Alexa Fluor 555 (cell wall). Images are average intensity projections of *z* stacks. Scale bars 2 μm. **b**, Cell volumes of SH1000 WT and *pbp3 pbp4* treated with 1x MIC MEM for 1 h as measured by fluorescence microscopy after NHS-ester Alexa Fluor 555 labelling (Fig. 3a). Each dot represents a single cell. The median of each distribution is indicated by a black line. The number of cells analysed for each condition was *n* ≥ 100. *P* value was determined by Mann–Whitney *U* tests (****, *P* < 0.0001). From left to right: *P* = 1.276e-042 and 1.421e- 024; *n* = 102, 100, 101 and 102. **c**, TEM of SH1000 WT and *pbp3 pbp4* treated with 1x MIC MEM for 1 h. Scale bars 200 nm. **d**, Quantification of phenotypes of SH1000 WT and *pbp3 pbp4* treated with MEM (1x MIC) for 1 h based on TEM data (Fig. 3c and Fig. 3 – figure supplement 1e, f). Examples of cells classified as normal (blue) are shown in Fig. 3 – figure supplement 1e, f. Cells with abnormal phenotypes (light blue) are shown in Fig. 3c and Fig. 3 – figure supplement 1e, f. From left to right *n*= 343, 287, 275 and 172. Data are representative of two (a, b (for WT) independent experiments. Experiments in b (for *pbp3 pbp4)* and d were performed once.

### PASTA domains mediate PBP1 interaction with division components

The morphologies of the *Δpbp1* and *pbp1_Δ_*_PASTA_ mutants resemble *S. aureus* depleted of DivIB in which EzrA and FtsZ form multiple rings and the synthesis of the cross wall is blocked, despite the normal recruitment of early cell division proteins and piecrust formation (Bottomley et al., 2014). In the *Δpbp1 ezrA-gfp* mutant, EzrA, which here acts as an early cell division marker, was localised at midcell in the majority of cells and formed additional arcs or rings in 33.5% of the population (Fig. 4a, d). Multiple EzrA rings were observed in 42.7% of the *pbp1_Δ_*_PASTA_ *ezrA-gfp* mutant cells (Fig. 4b, d), supporting the requirement for PBP1 PASTA domains for correct selection of the division site. Alternatively, the multiple division rings could result from a lack of the septal progression whereby the unproductive division machinery results in futile additional alternative initiation attempts, suggesting that PASTA domains are involved in the progression from piecrust to septal plate formation. While the number of cells with complete septa (EzrA-GFP visible as a line or focus) reduced by at least 6-fold in *Δpbp1 ezrA-gfp* and *pbp1_Δ_*_PASTA_ *ezrA-gfp*, it only halved in *pbp1* ezrA-gfp* (12.5% to 6.3% of *pbp1* ezrA-gfp* in +/-IPTG, respectively; Fig. 4c, d), confirming that septum progression, although reduced, still occurred when PBP1 TP was inactive implying that TP activity is necessary for correct septal architecture during cell division.

**Fig. 4.**
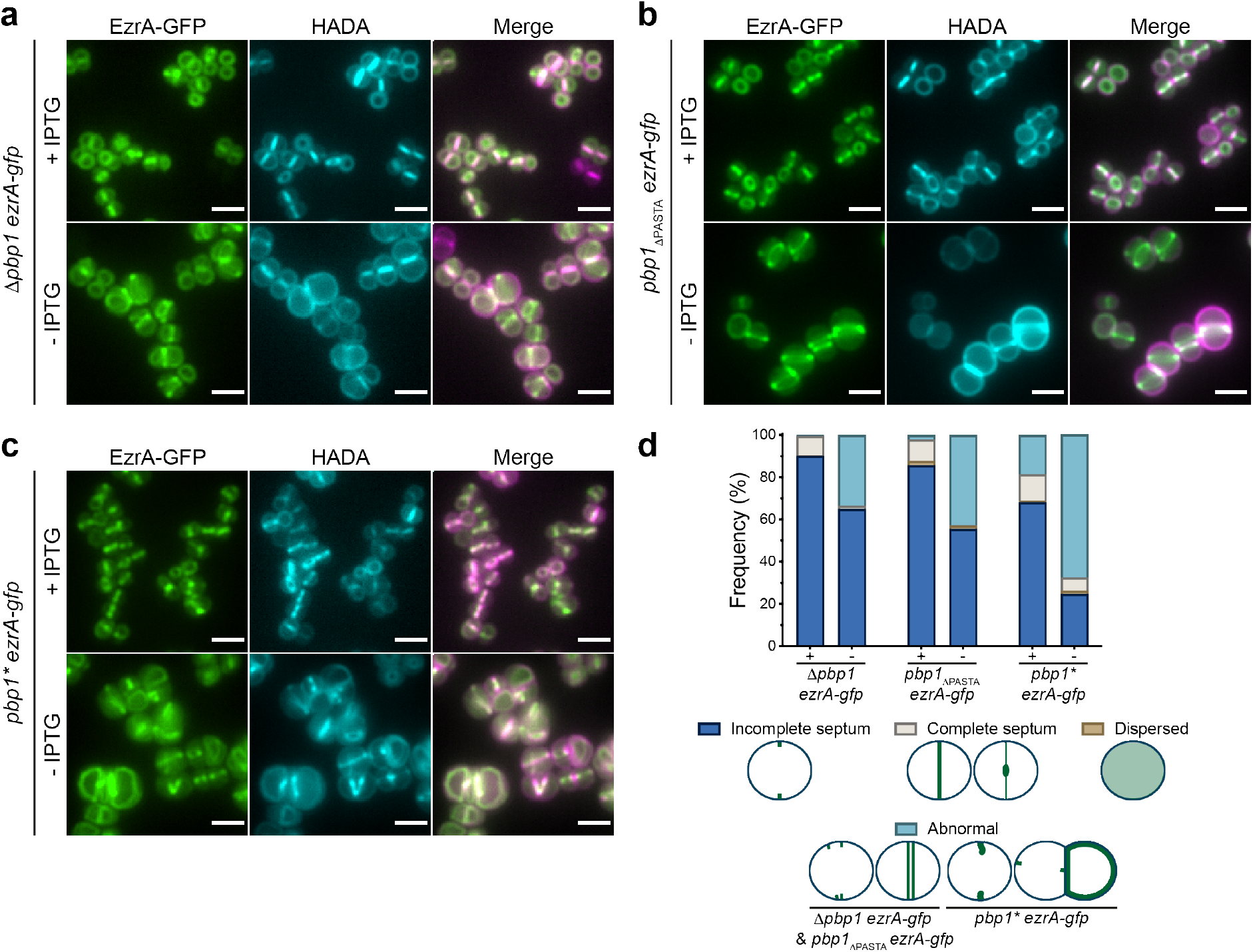
The role of PBP1, PASTA and TP domains in EzrA localisation in *S. aureus*. **a-c**, Localisation of EzrA-GFP in Δ*pbp1 ezrA-gfp, pbp1*_ΔPASTA_ *ezrA-gfp* and *pbp1* ezrA-gfp* grown in the presence or absence of IPTG for 2 h and labelled with HADA for 5 min to stain PG. Images are average intensity projections of *z* stacks. Scale bars 5 μm. **d**, Quantification of EzrA-GFP localisations in Δ*pbp1 ezrA-gfp*, *pbp1*_ΔPASTA_ *ezrA-gfp* and *pbp1* ezrA-gfp* grown with or without IPTG. From left to right *n =* 395, 499, 481, 438, 360 and 382. Data are representative of two independent experiments.

The cell wall of Gram-positive bacteria is decorated with wall teichoic acid (WTA) glycopolymers (Neuhaus & Baddiley, 2003). WTA regulates cell shape, ion homeostasis, autolytic enzymes, growth and division (Swoboda et al., 2010). In *S. aureus*, WTA plays a crucial role in virulence, MRSA resistance to *β*-lactam antibiotics, PBP4 localisation at the septum and PG cross-linking (Atilano et al., 2010; Campbell et al., 2011; Farha et al., 2013; Weidenmaier et al., 2005). Loss of WTA also results in a proportion of cells with aberrant septa (Campbell et al., 2011) suggesting a link with PBP1 function. Loss of *tarO* (leading to a lack of WTA) caused minor cell division defects in SH1000 (Fig. 4 – figure supplement 1a, e, f). Combining *tarO* with the mutations in *pbp1* exacerbated the observed morphological defects, with the appearance of distinct septal and off-septal PG foci appeared (marked with HADA) in *Δpbp1 tarO* and *pbp1_Δ_*_PASTA_ *tarO* (Fig. 4 – figure supplement 1b-f), demonstrating that both WTA and PBP1 are involved in cell cycle control in parallel.

As PBP1 PASTA has a role in the regulation of septal plate formation, this may be determined by interacting with other protein components. In order to examine this hypothesis we performed a bacterial two-hybrid assay, in which PBP1 has previously been found to have multiple interactions (Steele et al., 2011). Truncation of the PASTA domains not only reduced *S. aureus* PBP1 interaction with DivIB but also with FtsW, whilst recognition of other known interacting partners of PBP1 (EzrA, PBP2 and DivIC) were unaffected by the PASTA truncation (Fig. 5 – figure supplement 1a, b), suggesting that these wider interactions involve the N-terminal domain of PBP1.

### PBP1 PASTA domains bind peptidoglycan

Impaired interaction with DivIB could be one explanation for why cells depleted of PBP1 PASTA domains initiate irregular piecrusts and septation defects accrue as a consequence. PASTA domains have long been associated with PG binding because of work performed mainly on serine/threonine protein kinases (STPK) (Mir et al., 2011; Shah et al., 2008; Squeglia et al., 2011; Yeats et al., 2002). Therefore, we assessed whether *S. aureus* PBP1 and its PASTA domains could recognise PG by measuring their affinities for *S. aureus* cell wall PG with or without WTA (+/-WTA) with a semi-quantitative fluorescence binding assay (Bottomley et al., 2014) and *S. aureus* PBP1 derivatives produced in *Escherichia coli* (Fig. 5a and Fig. 5 – figure supplement 1c). Both *Sa*PBP1 (*K_d_* 19 ± 4 nM (+WTA), 115 ± 21 nM (- WTA)) and its PASTA domains (*Sa*PASTA_PBP1_; *K_d_* 198 ± 42 nM (+WTA), 109 ± 23 nM (-WTA) bound PG (Fig. 5b). Inactive *Sa*PBP1* was still able to bind PG with a preference for PG with WTA present (*K_d_* 53 ± 8 nM (+WTA), 227 ± 46 nM (-WTA); Fig. 5b), similar to active *Sa*PBP1. Although removal of the PASTA domains did not abolish BocillinFL binding (Fig. 5 – figure supplement 1c), it considerably reduced the ability of *Sa*PBP1*_Δ_*_PASTA_ to bind PG and binding was completely abolished in the presence of WTA (*K_d_* >2000 nM (+WTA), 440 ± 57 nM (-WTA); Fig. 5b). By contrast, the PASTA domains (*Sa*PASTA_PBP1_) on their own bind to *S. aureus* PG but are incapable of binding BocillinFL (Fig. 5b and Fig. 5 – figure supplement 1c). These results demonstrate unequivocally that PBP1 is a PG binding protein, and the PASTA domains have a dominant role in this interaction. Sequence conservation analysis of PASTA domains revealed the presence of either Arg or Glu residues in classifying a PASTA domain as a PG-binder (Calvanese et al., 2017). The PASTA domains of *S. aureus* PBP1 each have proline at the equivalent positions (residues Pro603 and Pro661) and thus PBP1 would be predicted as a non-PG binder, which clearly is not the case from the experimental evidence presented herein. Not only have we demonstrated the existence of such an interaction but we have also quantified it, suggesting that the predicted significance of conserved Arg or Glu residues with regard to PG binding is either only relevant to PASTA domains found in STPKs, linear arrangements of tandem PASTA repeats, or is too simplistic a prediction for proteins with multiple and complex functions like PBPs.

**Fig. 5.**
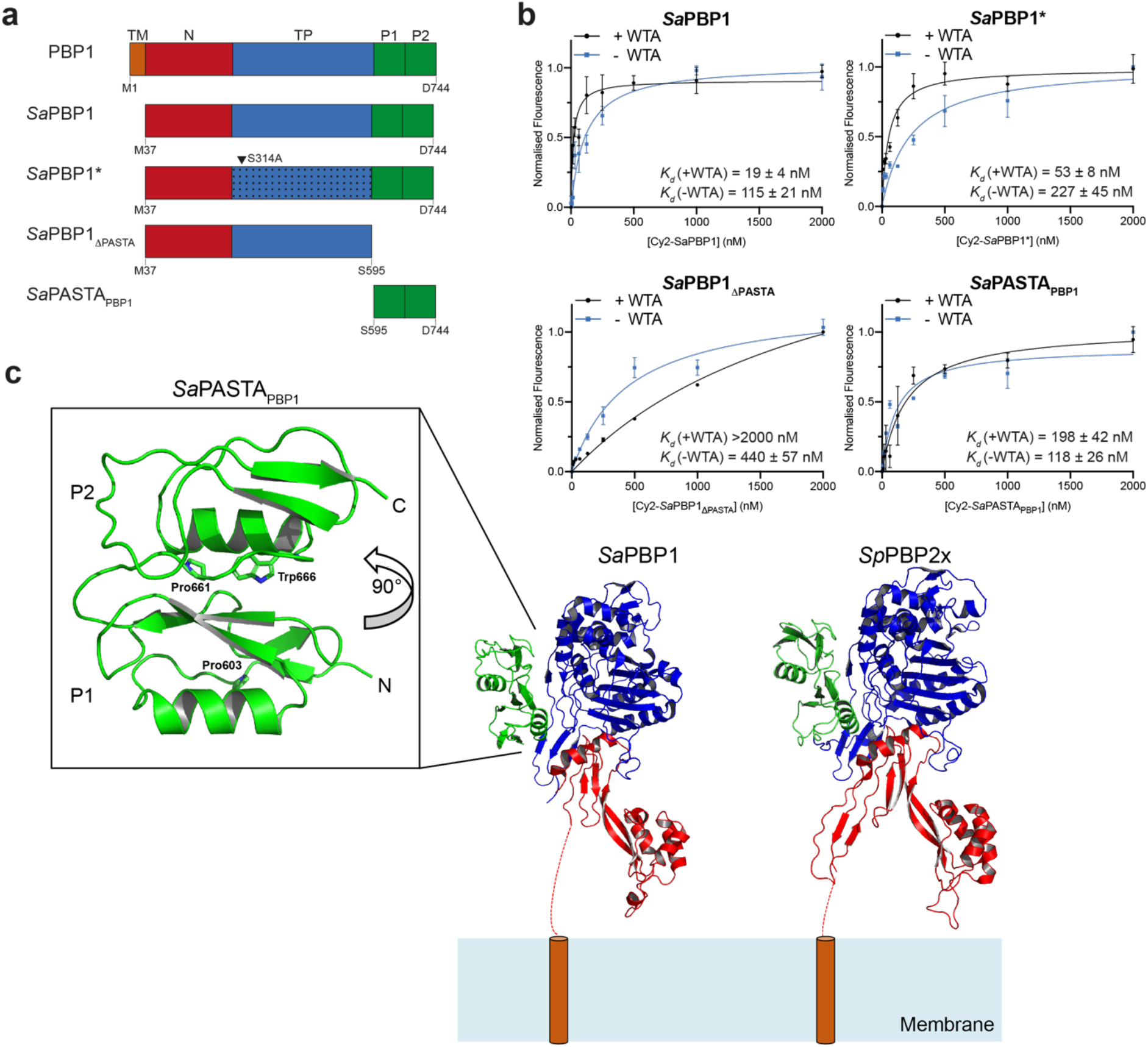
Role of PBP1 PASTA domains in cell wall binding. **a**, Schematic representation of structural domain organization of *S. aureus* PBP1 (top) and recombinant proteins (*Sa*PBP1, *Sa*PBP1_ΔPASTA_, *Sa*PBP1* and *Sa*PASTA_PBP1_) used in this study. TM, trans-membrane helix (orange); N, N-terminal dimerization domain (red); TP, transpeptidase domain (blue); P, PASTA domains (green). The arrowhead indicates the inactivation substitution in the TP domain of *Sa*PBP1*. The first and last amino acids of constructs are indicated. **b**, Fluorescent cell wall sedimentation assay. A Wilcoxon signed rank test (*P* < 0.05) was carried out to assess the significance of difference in +WTA/-WTA PG binding: *Sa*PBP1, *P* = 0.0273; *Sa*PBP1*, *P* = 0.0078; *Sa*PBP1_ΔPASTA_, *P* = 0.0195; *Sa*PASTA_PBP1_, *P* ≥ 0.9999, not significant. Data represent mean ± SD. Error bars that are smaller than the symbols are not shown. **c**, Structure of *Sa*PASTA_PBP1_. The crystal structure of *S. aureus* PBP1 lacking the PASTA domains (*Sa*PBP1, PDB 5TRO) and our *Sa*PASTA_PBP1_ structure were superimposed on to *S. pneumoniae* PBP2x (*Sp*PBP2x, PDB 5OAU) and displayed as cartoons. Their N-termini are orientated close to the representative cell membrane, as if anchored there by their respective transmembrane helices (dashed red line/orange cylinder). Individual domains are coloured as follows; N-terminal domain (red), transpeptidase domain (blue) and PASTA domain (green). The individual PASTA domains are labelled P1 and P2, respectively. Residues Pro603, Pro661 and Trp666 are displayed as sticks, with nitrogen atoms coloured blue.

To gain a better understanding of the role of the PASTA domains in *S. aureus* PBP1 (*Sa*PASTA_PBP1_), we determined their structure by X-ray crystallography. Soluble recombinant protein was obtained in high yield from the cytoplasm of *E. coli* cells and well- ordered crystals were subsequently produced that diffracted to a maximum resolution of 1.78 Å (Appendix Table 4). The structure was solved by molecular replacement using the corresponding PASTA domains present in *Sp*PBP2x from PDB entry 5OAU (Bernardo-García et al., 2018), which shares 26% sequence identity with *Sa*PASTA_PBP1_. The asymmetric unit contains two monomers (labelled A and B), each forming a 2-layer sandwich comprising an α-helix and a three-stranded antiparallel β-sheet, distinct from the TP domain (Fig. 5c).

Clear and continuous electron density allowed the modelling and unambiguous assignment of both PASTA domains (Fig. 5c). When *Sa*PASTA_PBP1_ is compared with other structures deposited in the PDB using *DALI* (Holm, 2020), the top hit identified was *S. pneumoniae* PBP2x (Z-score; 15.7), showing a significant conservation of the PASTA fold, despite low sequence identity (Fig. 5c). Unlike the linear arrangement observed for PASTA domains in serine/threonine kinases (Barthe et al., 2010; Ruggiero et al., 2011), *Sa*PASTA_PBP1_ adopts a compact upside-down globular arrangement (Fig. 5c). The arrangement of the two PASTA domains solved here, in isolation from the TP domain in comparison to structural analyses of *Sp*PBP2x, is entirely consistent with a non-linear PASTA domain arrangement. First, the structures of *Sa*PASTA_PBP1_ and the PASTA domains of *Sp*PBP2x share a pairwise RMSD of 2.2 Å over 114 C*α* and when *Sa*PASTA_PBP1_ is superimposed on the PASTA domains of *Sp*PBP2x there are no steric clashes with the TP domain. Second, the linker between PASTAs in *Sa*PASTA_PBP1_ has a sequence of DGDLTMPDMSGW, is neither glycine- nor alanine-rich, is not predicted to be disordered using IUPred2 or ANCHOR2 webservers and has a mean B factor of 44 Å^2^ in comparison to a mean B factor of 42 Å^2^ for the entire chain. Third, the interface between the PASTA domains is more reminiscent of the hydrophobic core of a globular protein than the more polar interface observed between molecules in crystal packing. It would therefore seem unlikely that the two PASTA domains open and close in a hinged manner, akin to the movement of a butterfly’s wings, in the presence and absence of endogenous PG. Finally, the two proline residues that apparently define PBP1 as a non-binder of PG are found buried from solvent either at the interface of PASTA domain 1 with the TP domain (Pro603) or at the interface between the TP domain and PASTA domains 1 and 2 (Pro661). This latter interface includes the only tryptophan (Trp666) in the sequence of *Sa*PASTA_PBP1_; tryptophan residues are frequent ‘markers’ of carbohydrate binding sites in proteins (Hudson et al., 2015) and in the absence of any obvious grooves or surface features associated with conserved sequence distributions and/or electrostatics it remains unclear how the PASTA domains of *Sa*PBP1 recognise PG.

## Discussion

*S. aureus* has just two essential PBPs (Reed et al., 2015) and so forms an apparently simple system to understand cell wall growth and division. Even the transpeptidase activity of these two enzymes can be substituted by a single enzyme in the presence of *β*-lactam antibiotics via the acquisition of PBP2A, encoded by *mecA*, in MRSA strains. Our recent study has revealed that the presence of *mecA* and associated genetic lesions have a profound effect on *S. aureus,* even in the absence of antibiotics (Panchal et al., 2020), leading to the discovery herein that the PG biosynthetic activity of PBP1 is essential in MSSA but not in MRSA (Fig. 1d). This observation has important ramifications for many studies in *S. aureus* where the use of an MRSA background can complicate phenotype interpretation. To understand the fundamental role of PBP1 activity in basic cell physiology we have thus used a MSSA strain with a defined genetic background.

The essential function of PBP1 is associated with its crucial role in septal PG synthesis (Pereira et al., 2009; Reichmann et al., 2019). Here we show that PBP1 is a multifunction regulatory and PG synthetic protein involved in both early and later stages of septum synthesis. PBP1 can interact with other cell division components, make and bind to PG. PG binding is primarily mediated by the PASTA domains that are essential for cell division. There is clear overall structural similarity between *S. aureus* PBP1 and *S. pneumoniae* PBP2x PASTA domains in the way that the two tandem PASTA domains associate into an anti- parallel bundle (Fig. 5c); this is in marked contrast to the head-to-tail linear PASTA domain repeats more typically found in STPKs. The highly hydrophobic interface between the two PASTA domains means it is unlikely to open up like butterfly wings to bind to PG; similarly an extensive, linear interaction with PG, which is likely to occur with the head-to-tail PASTA domain arrangements seen in STPKs and that may require their dimerization (Barthe et al., 2010), does not occur in *Sa*PBP1. Despite the successful production of diffracting crystals of *Sa*PASTA_PBP1_ grown in the presence of PG fragments (including an *N*-acetylglucosamine:*N*- acetylmuramic acid disaccharide) none of the structures yielded electron density features consistent with the stable binding of PG fragments. There are several potential explanations, including a lack of affinity of PASTA domains for small PG fragments, unrepresentative of the sacculus of *S. aureus*; our sedimentation assay does not permit the analysis of the binding of PASTA domains to small, soluble PG precursors. Consequently, and in common with all other PASTA domain structural analyses, the molecular details of PG recognition by *Sa*PBP1 remain elusive.

*S. aureus* is a spheroid coccus that can divide successively in three orthogonal planes (Saraiva et al., 2020; Turner et al., 2010). Septation is first observed as the formation of a thick band of PG known as the piecrust (Turner et al., 2010). This then transitions to the production of the septal plate itself, an initially V-shaped structure with a narrower leading edge (Lund et al., 2018). After closure of the septal annulus, the now bowed septum fills out to yield the mature structure prior to septal scission. The septal plate has two distinct PG architectures with a ring-like pattern at its core, which is exposed upon scission, and a subsequently-synthesised fine mesh, akin to the rest of the peripheral cell wall (Pasquina-Lemonche et al., 2020). Loss of the entire PBP1, or just its PASTA domains, does not prevent piecrust formation but does result in multi- and/or off-centre piecrusts without the ability to produce the septal plate (Fig 2f). Thus, piecrust formation does not require PBP1 but is likely the result of the activity of the essential PBP2. PBP1 may regulate division site selection through PG cell wall recognition via its PASTA domains. Alternatively, as the division apparatus is unable to progress effectively to septal plate formation due to the lack of PBP1, this may lead to further rounds of initiation and piecrust formation. PBP1 has a clear role in septal plate formation where in the absence of PBP1 or the PASTA domains, cells form aberrantly shaped septa that do not close their annuli (Fig. 2a-e). In stark contrast, inactivation of PBP1 TP activity (*pbp1\**) does not stop inward septum progression as observed with loss of PBP1 or the PASTA domains, however, such septa are mis-shapen, curved and abnormally thick (Fig. 2a-e and Fig. 3). The use of the PBP1-specific antibiotic MEM at 1x MIC led to the similar morphology of thickened and mis-shapen septa. Two independent avenues of research both lead to the conclusion that PBP1 TP activity is essential and, whilst septum formation is disturbed, it is not entirely prevented. Therefore PBP1 retains its regulatory function(s) regardless of activity loss. As well as binding to the cell wall, PBP1 also interacts with multiple protein partners including EzrA, DivIB/C, PBP2 and FtsW (Fig. 5 – figure supplement 1a, b) (Steele et al., 2011; Reichmann et al., 2019). Recently, the PASTA domains from *B. subtilis* PBP2B were shown to regulate PBP2B interaction with DivIB (Morales Angeles et al., 2020). *S. aureus* DivIB is a PG binding protein essential for division, which depletion leads to septal plate formation loss (Bottomley et al., 2014; Steele et al., 2011),. Here the PBP1 PASTA domains were found to bind DivIB and FtsW, alluding to their essential role in cell division. FtsW is a SEDS protein, whose TG activity requires the presence of PBP1 (Taguchi et al., 2019). Bifunctional aPBPs (including PBP2) and bPBP-SEDS (including PBP1-FtsW) pairs share similar activities but the fact they coexist in many bacterial species implies there is a division of responsibilities between them.

Indeed, it has been proposed lately that bPBP-SEDS pairs likely lay the primary PG matrix, while aPBPs support the initial PG by modifying, filling in and adding PG to it (Cho et al., 2016; Straume et al., 2020). The *S. aureus* septal plate PG has two distinct architectures, a disordered mesh present on its cytoplasm facing side and a ring structure at its core, which is revealed after the cells have split (Pasquina-Lemonche et al., 2020; Turner et al., 2010) (Fig. 5 – figure supplement 2). Recent AFM analysis from *Staphylococcus warneri* also describes the distinct PG architectures during septation as piecrust and septal plate rings/mesh (Su et al., 2020). When sacculi are purified from *S. warneri*, the septum can split apart revealing the rings, even in septa that have not closed their annulus, showing that the rings are not a result of PG hydrolysis during cell scission. We hypothesise (Fig. 5 – figure supplement 2) that once the piecrust has been produced, PBP1 and FtsW use this as a foundation to initiate septal plate formation. Together they make the rings of material that become the core of the developing septum, providing the framework for PBP2 to make the bulk of the septal plate as a tight mesh alongside PBP4 and the insertion of WTA via the *tar* pathway. Loss of PBP1 TP activity in the presence of active PBP2 leads to the lack of the ring framework and aberrant, unproductive septum formation. The rings that form the centre of the developing septum also provide the cleavage plane during scission.

Cell division is a fundamental requirement for life. A central question to this in bacteria is how is the division septum synthesised and then split to yield two daughter cells whilst maintaining cellular integrity in the face of internal turgor? Here we have begun to answer this question by revealing the complex synthesis coordination mechanisms that allow this biological engineering feat to be accomplished.

## Materials and methods

### Bacterial growth conditions

Strains used in this study are listed in Appendix Table 1.

All *Staphylococcus aureus* strains were grown in tryptic soy broth (TSB) containing appropriate antibiotics at 37°C, unless otherwise indicated, with aeration.

All *Escherichia coli* strains, unless otherwise stated, were grown in Lysogeny broth (LB) containing appropriate antibiotics at temperatures ranging from 20°C to 37°C with aeration. For solid media 1.5% (w/v) agar was added.

When necessary, growth medium was supplemented with kanamycin (50 μg ml^-1^), tetracycline (1 μg ml^-1^), chloramphenicol (10 μg ml^-1^, *S. aureus*; 30 μg ml^-1^, *E. coli*), erythromycin (5 μg ml^-1^), spectinomycin (250 μg ml ml^-1^), ampicillin (100 μg ml^-1^), meropenem (0.4 μg ml^-1^, 1x MIC for SH1000 WT; 0.2 μg ml^-1^, 1x MIC for *pbp3 pbp4*), 5- bromo-4-chloro-3-indolyl β-d-thiogalactopyranoside (X-Gal; 80 μg ml^-1^, *S. aureus*; 40 μg ml^-1^, *E. coli*) or isopropyl β-d-thiogalactopyranoside (IPTG, 50 μM or 1 mM).

### Plasmid construction

Plasmids and oligos used in this study are listed in Appendix Table 2 and Appendix Table 3, respectively.

Plasmids were cloned using *E. coli* NEB5*α* following previously described methods (Gibson et al., 2009; Sambrook et al., 1989).

### pKB-Pspac-pbp1

A fragment containing RBS and coding region of *S. aureus pbp1* was PCR amplified from the genomic DNA of *S. aureus* SH1000 using pCQ-pbp1-F/-R primers and cloned into NheI and AscI cut pCQ11-FtsZ-SNAP by Gibson assembly, resulting in pCQ11-P*spac*-*pbp1*. Next the region containing P*spac*, RBS and *pbp1* was PCR amplified from pCQ11-P*spac*-*pbp1* using pKB-Pspac-pbp1-F/-R primers and cloned into BamHI and EcoRI cut pKASBAR by Gibson assembly giving pKB-P*spac*-*pbp1*.

### pMAD-Δ*pbp1*

Fragments encompassing 1 kb regions flanking upstream (from -980 bp upstream of *pbp1* to first 20 bp of *pbp1*) and downstream of (from 2214 bp of *pbp1* to 970 bp downstream of *pbp1*) *pbp1* were PCR amplified from *S. aureus* SH1000 genomic DNA using primer pairs pbp1-A/-B and pbp1-C/-D, respectively, and cloned into BamHI and EcoRI cut pMAD by Gibson assembly, creating a deletion vector pMAD-*Δpbp1*.

### pMAD-*pbp1*Δ_PASTA_

Fragments encompassing 1.5 kb regions flanking the region encoding *pbp1* PASTA domains (upstream, from 286 bp to 1785 bp of *pbp1*; downstream, from 2214 bp of *pbp1* to 970 bp downstream of *pbp1*) were PCR amplified from *S. aureus* SH1000 genomic DNA using pbp1-E/-F and pbp1-G/-H primers and cloned into BamHI and EcoRI cut pMAD by Gibson assembly, resulting in a deletion vector pMAD-*pbp1_Δ_*_PASTA_.

### pMAD-pbp1*

A ∼1.3 kb fragment covering an upstream region of the active site of *pbp1* (from -334 bp upstream of *pbp1* to first 950 bp of the *pbp1* coding sequence), and a ∼1.3 kb fragment comprising the 3′ fragment of *pbp1* (930-2235 bp region of *pbp1*) were PCR amplified from *S. aureus* SH1000 genomic DNA using primer pairs pbp1*5′-F/-R and pbp1*3′-F/-R, respectively. Primers pbp1*5′-R and pbp1*3′-F were designed to introduce a T to G point mutation resulting in a Ser314Ala substitution. The PCR products were ligated with pMAD cut with EcoRI and BamHI by Gibson assembly, resulting in pMAD-*pbp1\**.

### T25-PBP1Δ_PASTA_

A fragment encoding *S. aureus pbp1* without the PASTA domains (M1-S595) was PCR amplified from *S. aureus* SH1000 genomic DNA using T25-pbp1-F and T25-pbp1pasta-R and cloned into BamHI and EcoRI cut pKT25, resulting in T25-PBP1*_Δ_*_PASTA_.

### pVR plasmids

Full-length *pbp1* (M1-D744) was *E. coli* codon optimised, synthesised by GenScript, PCR amplified using VR47F/R and cloned into KpnI and HindIII cut pOPINRSF using In-Fusion cloning (Takara Bio), resulting in pVR01. Construction of pVR02 (*Sa*PBP1, M37-D744) and pVR06 (*Sa*PASTA_PBP1_, S595-D744) was performed using inverse PCR (iPCR) (Erster & Liscovitch, 2010), pVR01 as a template and primer pairs VR49F/VR49R and VR57F/VR49R, respectively. pVR03 (*Sa*PBP1*, M37-D744, S314A) and pVR04 (*Sa*PBP1*_Δ_*_PASTA_, M37-S595), were constructed by QuikChange Site-Directed Mutagenesis of pVR02 using VR51 and VR53, respectively.

### pSA50

In order to construct an overexpression plasmid for sPBP1-BAP, A51-D744 fragment of *E. coli* codon optimised *pbp1* was PCR amplified using primers OPPF20018F/OPPF20018R and cloned into KpnI and SfoI cut pOPINJB by In-Fusion cloning (Takara Bio). The resulting construct, pSA50 contains an N-terminal hexahistidine-tag fused to Glutathione-S-transferase followed by a Human Rhinovirus 3C protease site, while the PBP1 (A51-D744) C-terminal end is fused to a Biotin Acceptor Peptide (BAP) sequence.

### Construction of *S. aureus* mutants

All vectors were passed through a restriction-deficient *S. aureus* RN4220 before being transduced into a final *S. aureus* SH1000 strain. Transformation and phage transduction of *S. aureus* were carried out as described previously (Novick & Morse, 1967; Schenk & Laddaga, 1992).

### Δpbp1, pbp1Δ_PASTA_ and pbp1*

For construction of *pbp1* mutation strains, first an ectopic copy of *pbp1* under the control of the P*spac* promoter was introduced at the lipase (*geh*) locus. Electrocompetent CYL316 was transformed with pKB-P*spac*-*pbp1*. The chromosomal fragment containing the integrated plasmid was moved into *S. aureus* SH1000 by phage transduction, resulting in SJF4588 (*S. aureus* SH1000 *geh::*P*spac-pbp1*). Next electrocompetent RN4220 was transformed with pMAD-*Δpbp1*, pMAD-*pbp1_Δ_*_PASTA_ or pMAD-*pbp1\** and the plasmids were moved to SJF4588 by phage transduction. Integration at 42°C and excision at 28°C of pMAD-*Δpbp1*, pMAD-*pbp1_Δ_*_PASTA_ or pMAD-*pbp1\**, resulted in strains SJF5116, SJF5275 and SJF4590, respectively. To allow controlled expression of *pbp1* from P*spac*, pGL485, a multi-copy plasmid carrying *lacI*, was introduced creating strains *Δpbp1* (*S. aureus* SH1000 *geh::*P*spac- pbp1 Δpbp1 lacI*), *pbp1_Δ_*_PASTA_ (*S. aureus* SH1000 *geh::*P*spac-pbp1 pbp1_Δ_*_PASTA_ *lacI*) and *pbp1\** (*S. aureus* SH1000 *geh::*P*spac-pbp1 pbp1* lacI*).

### MRSA Δ*pbp1* and MRSA *pbp1\**

In order to construct high-level *β*-lactam resistant mutants, *Δpbp1* and *pbp1\** were transformed with a phage lysate from SJF5046 (*S. aureus* SH1000*lysA::*p*mecA rpoB*^H929Q^) with selection for erythromycin resistance, resulting in low-level *β*-lactam resistant *Δpbp1* p*mecA* and *pbp1\** p*mecA.* The low-level resistant mutants were transduced again with the phage lysate from SJF5046 and selected for kanamycin resistance, resulting in MRSA *Δpbp1* (*S. aureus* SH1000 *geh::*P*spac-pbp1 Δpbp1 lacI lysA::*p*mecA rpoB*^H929Q^) and MRSA *pbp1* pbp1* (*S. aureus* SH1000 *geh::*P*spac-pbp1 pbp1* lacI lysA::*p*mecA rpoB*^H929Q^). MIC values were determined using antibiotic susceptibility tests using E-test M.I.C. Evaluator (Oxoid) strips.

### pbp3 pbp4

SH1000 was transduced with a phage lysate from NE420 (*S. aureus* JE2 *pbp3::Tn*) resulting in SH4421 (*S. aureus* SH1000 *pbp3::Tn*). To swap the erythromycin resistance cassette to a kanamycin cassette, SH4425 (*S. aureus* SH1000 *pbp4::Tn*) was transduced with a phage lysate from NE3004 (*S. aureus* RN4220 pKAN). Integration at 42°C and excision at 28°C of pSPC resulted in strain SH5115 (*S. aureus* SH1000 *pbp4::kan*). SH4421 was subsequently transduced with a phage lysate from SH5115 (*S. aureus* SH1000 *pbp4::Tn*) resulting in *pbp3 pbp4* (SH5483; *S. aureus* SH1000 *pbp3::Tn pbp4::kan*).

### Δpbp1 pbp4, pbp1Δ_PASTA_ pbp4 and pbp1* pbp4

*Δpbp1, pbp1_Δ_*_PASTA_ and *pbp1\** were transduced with a phage lysate from SH5115 (*S. aureus* SH1000 *pbp4::kan*), resulting in *Δpbp1 pbp4* (*S. aureus* SH1000 *geh::*P*spac-pbp1 Δpbp1 lacI pbp4::kan*)*, pbp1_Δ_*_PASTA_ *pbp4* (*S. aureus* SH1000 *geh::*P*spac-pbp1 pbp1_Δ_*_PASTA_ *lacI pbp4::kan*) and *pbp1* pbp4* (*S. aureus* SH1000 *geh::*P*spac-pbp1 pbp1* lacI pbp4::kan*), respectively.

### pbp1* pbp3 pbp4

*pbp1* pbp4* (*S. aureus* SH1000 *geh::*P*spac-pbp1 pbp1* lacI pbp4::kan*) was transduced with a phage lysate from SH4421 (*S. aureus* SH1000 *pbp3::Tn*), resulting in *pbp1* pbp3 pbp4* (*S. aureus* SH1000 *geh::*P*spac-pbp1 pbp1* lacI pbp3::Tn pbp4::kan*).

### Δpbp1 ezrA-gfp, pbp1Δ_PASTA_ ezrA-gfp and pbp1* ezrA-gfp

*Δpbp1, pbp1_Δ_*_PASTA_ and *pbp1\** were transduced with a phage lysate from JGL227 (*S. aureus* SH1000 *ezrA-gfp+*), resulting in *Δpbp1 ezrA-gfp* (*S. aureus* SH1000 *geh::*P*spac-pbp1 Δpbp1 lacI ezrA-gfp*)*, pbp1_Δ_*_PASTA_ *ezrA-gfp* _PASTA_ (*S. aureus* SH1000 *geh::*P*spac-pbp1 pbp1_Δ_*_PASTA_ *lacI ezrA-gfp*) and *pbp1* ezrA-gfp* (*S. aureus* SH1000 *geh::*P*spac-pbp1 pbp1* lacI ezrA-gfp*), respectively.

### Δpbp1 tarO, pbp1Δ_PASTA_ tarO and pbp1* tarO

*Δpbp1, pbp1_Δ_*_PASTA_ and *pbp1\** were transduced with a phage lysate from *tarO tarO+* (*S. aureus* SA113 *ΔtarO::ery* pUC1-*tarO*), resulting in *Δpbp1 tarO* (*S. aureus* SH1000 *geh::*P*spac-pbp1 Δpbp1 lacI ΔtarO::ery*)*, pbp1_Δ_*_PASTA_ *tarO* _PASTA_ (*S. aureus* SH1000 *geh::*P*spac-pbp1 pbp1_Δ_*_PASTA_ *lacI ΔtarO::ery*) and *pbp1* tarO* (*S. aureus* SH1000 *geh::*P*spac-pbp1 pbp1* lacI ΔtarO::ery*), respectively.

### PBP1 depletion

P*spac-pbp1* strains were grown from an OD_600_ of 0.1 to exponential phase (OD_600_ ∼ 0.5) in TSB containing 10 µg ml^−1^ chloramphenicol and 50 µM IPTG. Cells were washed three times by centrifugation and resuspension in TSB. Washed cells were then used to inoculate TSB 10 µg ml^−1^ chloramphenicol. Cultures were inoculated to an OD_600_ 0.05 for phenotypic studies and an OD_600_ 0.005 for growth studies. For phenotypic analysis, cultures were incubated for 2 h to allow depletion of PBP1 before microscopy imaging. Control samples were grown in TSB supplemented with 10 µg ml^−1^ chloramphenicol and 1 mM (50 µM, *ezrA-gfp* mutants) IPTG.

For the plating efficiency test, cells grown in the presence of 10 µg ml^−1^ chloramphenicol and 50 µM IPTG to exponential phase (OD_600_ ∼ 0.5) were washed three times in PBS. Serial dilutions of washed cells were plated on TSB 10 µg ml^−1^ chloramphenicol with or without 1 mM IPTG. Relative plating efficiency (%*CFU*) is expressed as the number of cells that grow on plates without IPTG (*CFU*_%& ()*+_) to cells that grow in the presence of IPTG (*CFU*_()*+_) multiplied by 100%:

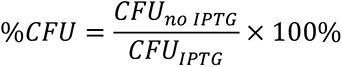

### Meropenem activity assays

*S. aureus* strains were grown overnight in TSB. The overnight cultures were used to inoculate fresh TSB media to an OD_600_ of 0.05. When cells reached an OD_600_ of 0.2-0.4, meropenem was added, and the change of bacterial count was monitored. The colony-forming units per ml of culture (CFU/ml) measures were normalized to the initial CFU/ml at the time of the antibiotic addition, at time zero (*t_0_*).

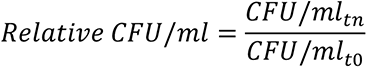

For phenotypic analysis, cells were treated for 1 h with 1x MIC meropenem before microscopy imaging.

### Fractionation of *S. aureus* membranes

The membrane fraction of *S. aureus* was prepared as previously described (García-Lara et al., 2015) with the following modifications. *S. aureus* cells grown to the appropriate growth phase were recovered by centrifugation (5,000 x g, 10 min, 4°C), washed three times by resuspension and centrifugation (5,000 x g, 10 min, 4°C) in PBS. Cells were resuspended in TBSI and broken using 0.1 mm silica spheres (Lysing Matrix B) and FastPrep Homogenizer (MP Biomedicals) in 12 cycles of 30 s, at maximum speed (6.5 m s^-1^), with 5 min incubation on ice between cycles. Cell lysates were centrifuged (5,000 x g, 10 min, 4°C) to remove unbroken cells. The supernatant was then spun (5,000 x g, 10 min, 4°C) to sediment cell wall material. The membrane fraction was recovered from the supernatant by centrifugation (35,000 x g, 20 min, 4°C) and the pellet (membranes) was resuspended in PBS. The total protein concentration was estimated by Bradford assay.

### *In vitro* labelling of *S. aureus* PBPs with BocillinFL

This method was adopted from a published protocol (Zhao et al., 1999) with minor modifications. Membrane proteome samples (25 μg in 20 μl PBS) and purified proteins (2.5 μg in 25 μl HEPES pH 7.5 150 mM NaCl) were incubated with 25 μM BocillinFL (Invitrogen) for 20 min at 37°C. Additionally for competition assay, purified *Sa*PBP1 was mixed with 2.5 µg (∼286 µM final concentration) ampicillin and incubated at 37°C for 10 min prior to the addition of BocillinFL. The reaction was stopped by the addition of 5x SDS-PAGE loading buffer. Membrane proteome was additionally incubated for 10 min at 90°C. The samples were run on a 6-20% (w/v) SDS-PAGE gradient or 10% (w/v) SDS-PAGE gel and visualized using a BioRad ChemiDoc MP Imaging system or a GE Typhoon FLA 9500.

### Labelling *S. aureus* DAAs

*S. aureus* cells were incubated with 500 µM (2 mM for *pbp4* mutants) HADA or 1 mM ADA-DA at 37°C for 5 min. Cells were then washed by centrifugation and resuspension in PBS.

### Click chemistry

ADA-DA containing an azide functional group was fluorescently labelled with Alexa Fluor 488 Alkyne at 5 µg ml^−1^ via the Click reaction (copper (I)-catalysed alkyne-azide cycloaddition). This was carried out using the Click-iT Cell Reaction Buffer Kit (ThermoFisher) according to the manufacturer’s protocol.

### Labelling *S. aureus* with fluorescent NHS-ester

Fixed cells wells were resuspended in PBS containing 8 μg ml^−1^ Alexa Fluor 555 NHS ester (Invitrogen) and incubated at room temperature for 30 min. Cells were washed twice by centrifugation and resuspension in PBS.

### Fixing for fluorescence microscopy

Cells were fixed by incubation in 1.6% (w/v) paraformaldehyde at room temperature for 30 min.

### Fluorescence microscopy

Fixed cells were dried onto a poly-l-Lysine coated slide, mounted in PBPS and imaged on a Nikon Ti Inverted microscope fitted with a Lumencor Spectra X light engine. Images were taken using a 100x PlanApo (1.4 NA) oil objective using 1.518 RI oil and detected by an Andor Zyla sCMOS camera.

### Cell volume estimation

Cell volume calculations were carried out as previously described (Zhou et al., 2015). The long and short axis of cells were measured using Fiji. The volume was then calculated based on a prolate spheroid shape with volume:

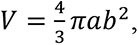

where *a* and *b* are the radii along the long and short axis, respectively.

### Transmission electron microscopy

*S. aureus* strains were prepared for electron microscopy as previously described (Sutton et al., 2021).

### Preparation of *S. aureus* sacculi

Peptidoglycan from *S. aureus* cells was extracted and if required HF treated to remove cell wall polymers as previously described (Sutton et al., 2021).

### Sacculi immobilisation for AFM Imaging

Immobilisation surface was prepared by adding the solution mixed by 171 µL 100 mM NaHCO_3_, 3 µl of 1 M NaOH and 6 µl of Cell-Tak (Corning, 5% (w/v) in acetic acid) on freshly cleaved mica. After 30 min incubation, the surface was washed by 5 × 200 µl HPLC grade water. Sacculi stocks were 10 times diluted in HPLC grade water and briefly tip- sonicated to re-suspend in prior to immobilisation. 10 µl of the sacculi suspension was added to 40 µl of HPLC grade water on the Cell-Tak immobilisation surface and incubated for 1 h. The surface was then thoroughly rinsed with HPLC grade water, blow-dried with nitrogen and stored in petri-dish at room temperature before AFM imaging.

### AFM Imaging and image analysis

AFM imaging was carried out on a Nanowizard III ULTRA Speed system (JPK, Germany). Rectangular cantilevers with nominal spring constant of 0.3 N/m and resonant frequency (in liquid) of ∼ 150 kHz (USC-F0.3-k0.3, NanoWorld, Switzerland) were used. The spring constant and deflection sensitivity of each cantilever was calibrated prior to each measurement (Hutter & Bechhoefer, 1993; Sader et al., 2016) Measurements were carried out in Quantitative Imaging mode at room temperature, in the buffer composed of 200 mM KCl and 10 mM Tris. Scans were driven at a line rate of ∼ 0.78 Hz, with a typical Z length of 300 nm and trigger force of 20 nN.

Resultant topographic images were processed using JPK Data Processing. No flattening or surface subtraction was applied. High pass filter (scale: 100% to 500%, degree of smoothing: 5 px, horizontal) was applied to the higher magnification images to enhance the contrast without modifying the morphological features. The morphological features of sacculi were summarised from images obtained on abundant technical repeats of 2 biological replicates.

### Recombinant protein production and purification sPBP1-BAP

*E. coli* BL21(DE3) cells containing plasmid pSA50 were grown in LB medium supplemented with 100 µg ml^-1^ ampicillin at 37°C to an OD_578_ of 0.5. Protein overproduction was induced by addition of 0.5 mM IPTG to the cell culture and further incubation for 4 h at 30°C. Cells were harvested by centrifugation (6,200 x g, 15 min, 4°C) and the pellet was resuspended in basic buffer (25 mM Tris-HCl, 100 mM NaCl, pH 7.5). After addition of 1 mM PMSF, 1:1,000 dilution of protease inhibitor cocktail (Sigma-Aldrich) and DNase, the cells were disrupted by sonication (Branson Digital Sonifier). The cell lysate was centrifuged (130,000 x g, 60 min, 4°C) and the supernatant was recovered. The supernatant was incubated with Ni-NTA Superflow (Qiagen) for 2 h at 4°C with gentle stirring, which had been pre-equilibrated in basic buffer. The resin was poured into a gravity column and washed with 20 volumes of wash buffer (25 mM Tris-HCl, 150 mM NaCl, 10% (v/v) glycerol, 10 mM MgCl_2_, 20 mM imidazole, pH 7.5). Bound protein was eluted with elution buffer (25 mM Tris-HCl, 150 mM NaCl, 10% (v/v) glycerol, 10 mM MgCl_2_, 600 mM imidazole, pH 7.5). 10 U ml−1 of HRV- 3C protease (Takara) were added to the Ni-NTA eluted protein to remove the oligohistidine- GST-tag during dialysis against 3 L of dialysis buffer I (25 mM Tris-HCl, 150 mM NaCl, 10 mM EGTA, 10% (v/v) glycerol, pH 7.5) for 20 h at 4°C. Digested protein was dialysed against 3 L of dialysis buffer II (25 mM Tris-HCl, 150 mM NaCl, 10 mM MgCl_2_, 10% (v/v) glycerol, pH 7.5), for 3 h at 4°C. The protein was incubated in the same Ni-NTA beads (pre-equilibrated in dialysis buffer II) for 2 h at 4°C to remove the contaminants and the His-GST tag from the sample. The flow through and the washes (2 volume of wash buffer) were pooled, dialysed against storage buffer (25 mM HEPES-NaOH, 150 mM NaCl, 10 mM MgCl_2_, 10% (v/v) glycerol, pH 7.5) and concentrated using a Vivaspin Turbo 15 column (MWCO 50000 Da).

### *Sa*PBP1, *Sa*PBP1*, *Sa*PBP1Δ_PASTA_ and *Sa*PASTA_PBP1_

All recombinant proteins were produced in *E. coli* Rosetta (DE3) cells at 37°C in TB medium supplemented with 50 µg ml^-1^ kanamycin and 30 µg ml^-1^ chloramphenicol. Once cultures had reached OD_600_ 0.9, protein expression was induced with 1 mM IPTG for 20 h at 20°C. Cells were harvested by centrifugation (4,000 x g at 4°C for 30 mins) and the pellet was resuspended in a buffer of 50 mM Tris-HCl pH 8.0, 500 mM NaCl, 20 mM imidazole supplemented with one EDTA-free protease inhibitor cocktail tablet (Roche) and DNAse (4 µg ml^-1^ final concentration). Cells in this resuspension were lysed by two passes through a One-Shot cell disruptor (Constant Systems) at 23 kpsi and the cell debris was removed by centrifugation (40,000 x g at 4°C for 30 min). The first purification step was affinity chromatography with a 5 mL HisTrap™ FF column (GE Healthcare) precharged with Ni^2+^ and equilibrated in buffer A (50 mM Tris-HCl, pH 8.0, 500 mM NaCl, 20 mM imidazole). A linear concentration gradient of imidazole was applied to elute the protein using buffer B (50 mM Tris-HCl, pH 8.0, 500 mM NaCl, 800 mM imidazole). Further purification was carried out by size exclusion chromatography using a Superdex™ 200 Hi Load 16/60 column (GE Healthcare). Proteins were eluted with SEC buffer (25 mM Tris-HCl, pH 8.0, 150mM NaCl) and analysed by SDS-PAGE.

### Generation of anti-PBP1 antibody

Serum against sPBP1A-BAP was produced from rabbits following a 28-day immunisation program at Eurogentec (Belgium), and it was purified as previously described (Bertsche et al., 2006).

### Immunoblot analysis

*S. aureus* cultures were washed three times by resuspension and centrifugation (5,000 x g, 10 min, 4°C) in PBS. Cells were resuspended in TBSI (50 mM Tris, 100 mM NaCl, pH 8, plus Complete Protease Inhibitor Cocktail, Roche) and broken using 0.1 mm silica spheres (Lysing Matrix B) and FastPrep Homogenizer (MP Biomedicals) in 12 cycles of 30 s, at maximum speed (6.5 m s^-1^), with 5 min incubation on ice between cycles. Cell lysates were centrifuged (5,000 x g, 10 min, 4°C) to remove unbroken cells. ∼ 60 μg of total protein was separated on a 12% (w/v) SDS-PAGE gel and electroblotted onto a nitrocellulose membrane and blocked in 5% (w/v) skimmed-milk in TBST (20 mM Tris-HCl, pH 7.6; 17 mM NaCl, 0.1% (v/v) Tween-20). The membrane blocked in 5% (w/v) skimmed-milk in TBST (20 mM Tris-HCl, pH 7.6; 17 mM NaCl, 0.1% (v/v) Tween-20) was incubated with primary polyclonal anti-PBP1 (1:1,000) overnight with gentle agitation at 4°C. Primary antibodies were detected using horseradish peroxidase-conjugated goat anti-rabbit IgG (1:10,000, BioRad) and Clarity Western ECL Substrate (BioRad) reagent according to the manufacturer’s protocol. Chemiluminescence was detected using Syngene G:BOX Chemi XX9.

### Crystallisation, data collection and structure determination

Crystallisation of *Sa*PASTA_PBP1_ was carried out at 20°C by the sitting-drop vapour diffusion method in 96-well MRC plates (Molecular Dimensions) with a Mosquito crystallisation robot (TTP LabTech) and commercial crystallisation screens (Hampton Research and Molecular Dimensions). Orthorhombic crystals of diffraction quality, with a maximum dimension of approximately 500 µm, appeared overnight from a mixture of equal volumes of protein solution (42 mg ml^-1^ in 25 mM Tris-HCl, pH 8.0, 150 mM NaCl) and reservoir solution (0.2 M NaCl, 0.1 M sodium/potassium phosphate pH 6.2, 50 % (v/v) PEG 200). Diffraction data were indexed and integrated using XDS (Kabsch, 2010) and scaled using AIMLESS (Evans & Murshudov, 2013) from the CCP4 program suite (Winn et al., 2011). The crystals display space group *P*2_1_2_1_2_1_ with unit cell lengths *a* = 39.8 Å, *b* = 81.4 Å and *c* = 89.6 Å. The asymmetric unit consists of two polypeptide chains with an estimated solvent content of 45 % and a V_m_ of 2.24 Å^3^/Da. The region corresponding to the two PASTA domains in the crystal structure of *S. pneumoniae* PBP2x (PBP 5OAU) was used as a molecular replacement search model, sharing approximately 26 % sequence identity with *Sa*PASTA_PBP1_. The search model was generated using phenix.sculptor (Bunkóczi & Read, 2011) to remove non- macromolecular chains and prune sidechains. The structure was solved by molecular replacement using PHASER (McCoy et al., 2007) and the resultant electron density map was of high quality, allowing the tracing of the main chain. Model building and refinement were carried out with Coot (Emsley et al., 2010) and Phenix (Liebschner et al., 2019), respectively. Assessments of the geometry and validation of the final model was carried out using Molprobity (Chen et al., 2010). Analyses of surface areas and interactions were made using the PISA web service (Krissinel & Henrick, 2007). The graphics program PyMOL (Schrӧdinger, LLC, 2015) was used to generate all molecular figures presented.

### Cell wall binding assays

Cell wall binding assays of recombinant PBP1 proteins fluorescently labelled with Cy2 bis-reactive dye (GE Healthcare) were performed as previously described (Bottomley et al., 2014).

### Bacterial two-hybrid

Competent BTH101 was co-transformed with pKT25 and pUT18 derivatives. Transformants were selected on LB agar plates containing 100 μg ml^−1^ ampicillin, 50 μg ml^−1^ kanamycin and 40 μg ml^−1^ X-Gal and incubated at 30°C. Single colonies were grown in 150 μl LB with 100 μg ml^−1^ ampicillin, 50 μg ml^−1^ kanamycin and 0.5 mM IPTG at 30°C.

To qualitatively measure for pairwise interactions, 5 μl of each overnight culture were spotted onto LB agar plates containing 100 μg ml^−1^ ampicillin, 50 μg ml^−1^ kanamycin, 0.5 mM IPTG and 40 μg ml^−1^ X-Gal. Plates were incubated at 30°C 24 – 48 h in an environment protected from light and imaged. To quantify interactions, overnight cultures were assayed for β-galactosidase activity against MUG (4-methylumbelliferyl-β-d-galactopyranoside) using an assay as previously described (Steele et al., 2011).

## Acknowledgements

This work was funded by the Medical Research Council (MR/N002679/1; MR/K015753/1), UKRI Strategic Priorities Fund (EP/T002778/1) and the Wellcome Trust (212197/Z/19/Z). L.L. thanks The Florey Institute for her PhD studentship. We gratefully acknowledge the Wolfson Light Microscopy facility for their support and assistance in this work. We thank Diamond Light Source for access to beamline I24 (mx18598) and Arnaud Baslé for data collection, help with figure generation and support. We are grateful to Joshua Sutton, Grace Pidwill, Victoria Lund, Laia Pasquina-Lemonche and Christopher Hill for help and advice.

## Supplementary figures and legends

**Fig. 1 – figure supplement 1.**
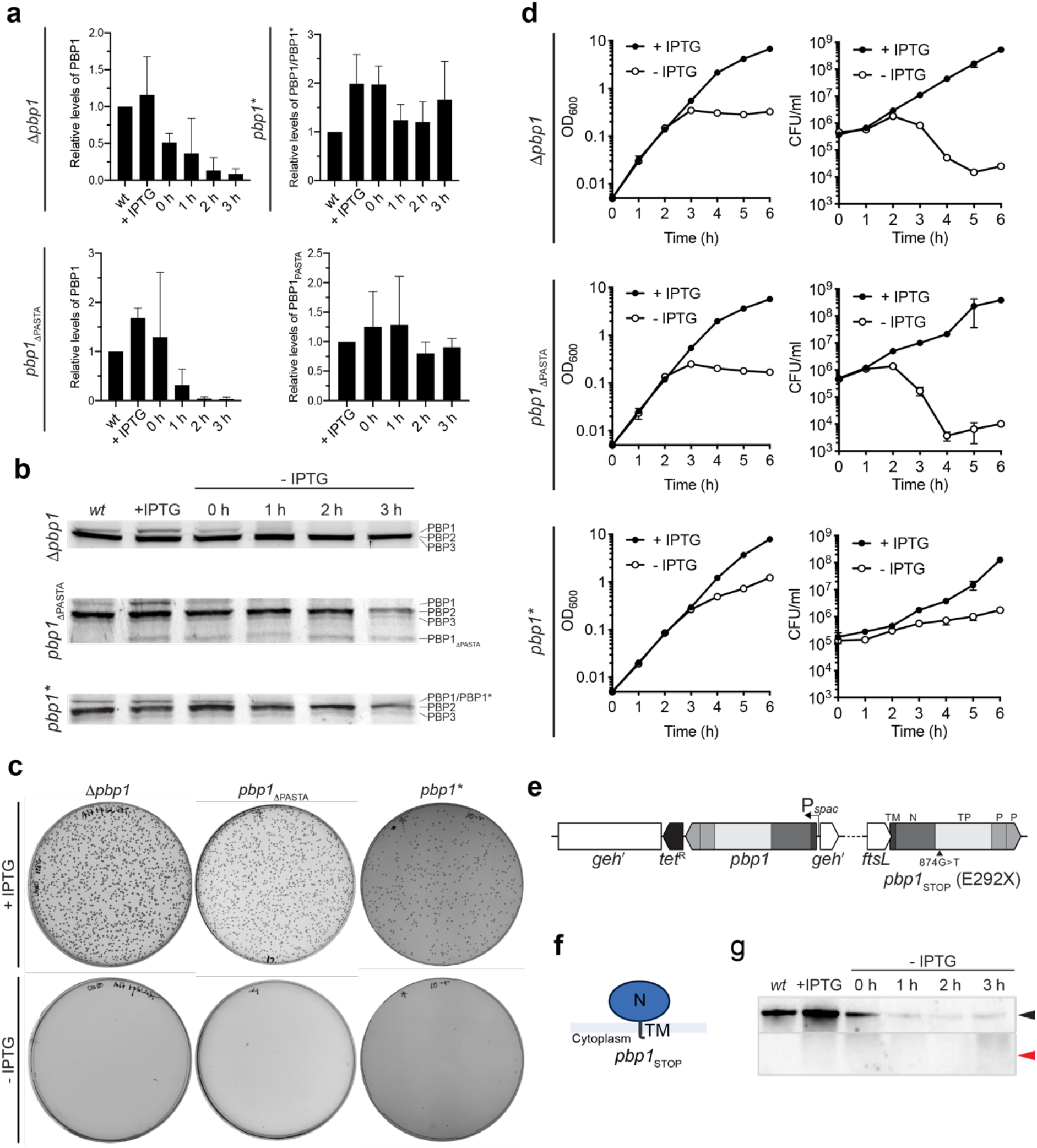
Essentiality of PBP1, PASTA and TP domains in *S. aureus.* **a**, Relative levels of PBP1 in Δ*pbp1, pbp1*_ΔPASTA_ and *pbp1\** grown with IPTG and for 0, 1, 2 and 3 h after inducer removal. PBP1 levels were normalized to protein levels in SH1000 *lacI* (*wt*). PBP1_ΔPASTA_ levels were normalized to PBP1_ΔPASTA_ levels in the presence of inducer. Quantifications are for the data shown in Fig. 1c. Data represent mean ± SD. **b**, BocillinFL gel-based analysis of penicillin binding proteins in SH1000 *lacI* (*wt*) and Δ*pbp1, pbp1*_ΔPASTA_ and *pbp1\** grown with IPTG and 0, 1, 2 and 3 h after inducer removal. **c**, Growth of Δ*pbp1, pbp1*_ΔPASTA_ and *pbp1\** with or without IPTG. Quantifications for the plate assay are shown in Fig. 1d. **d**, Growth curves of Δ*pbp1, pbp1*_ΔPASTA_ and *pbp1\** in the presence or absence of IPTG. Data represent mean ± SD. Error bars that are smaller than the data point symbols are not shown. **e**, Schematic representation of the *pbp1*_STOP_ mutant. A SNP (874 G for T) resulted in a premature stop codon (E292X) and removal of the TP and PASTA domains. **f**, Schematic representation of domain architecture of PBP1_STOP_ encoded by the *pbp1*_STOP_ mutant. **g**, Immunoblot showing PBP1 levels in SH1000 *lacI* (*wt*) and *pbp1*_STOP_ grown with IPTG and for 0, 1, 2 and 3 h without inducer analysed using anti-PBP1 antibody. Expected sizes: PBP1 = 83 kDa (black arrowhead) and PBP1_STOP_ = 33 kDa (red arrowhead). Data are representative of two (g), three (a, b, d) and at least four (c) independent experiments.

**Fig. 1 – figure supplement 2.**
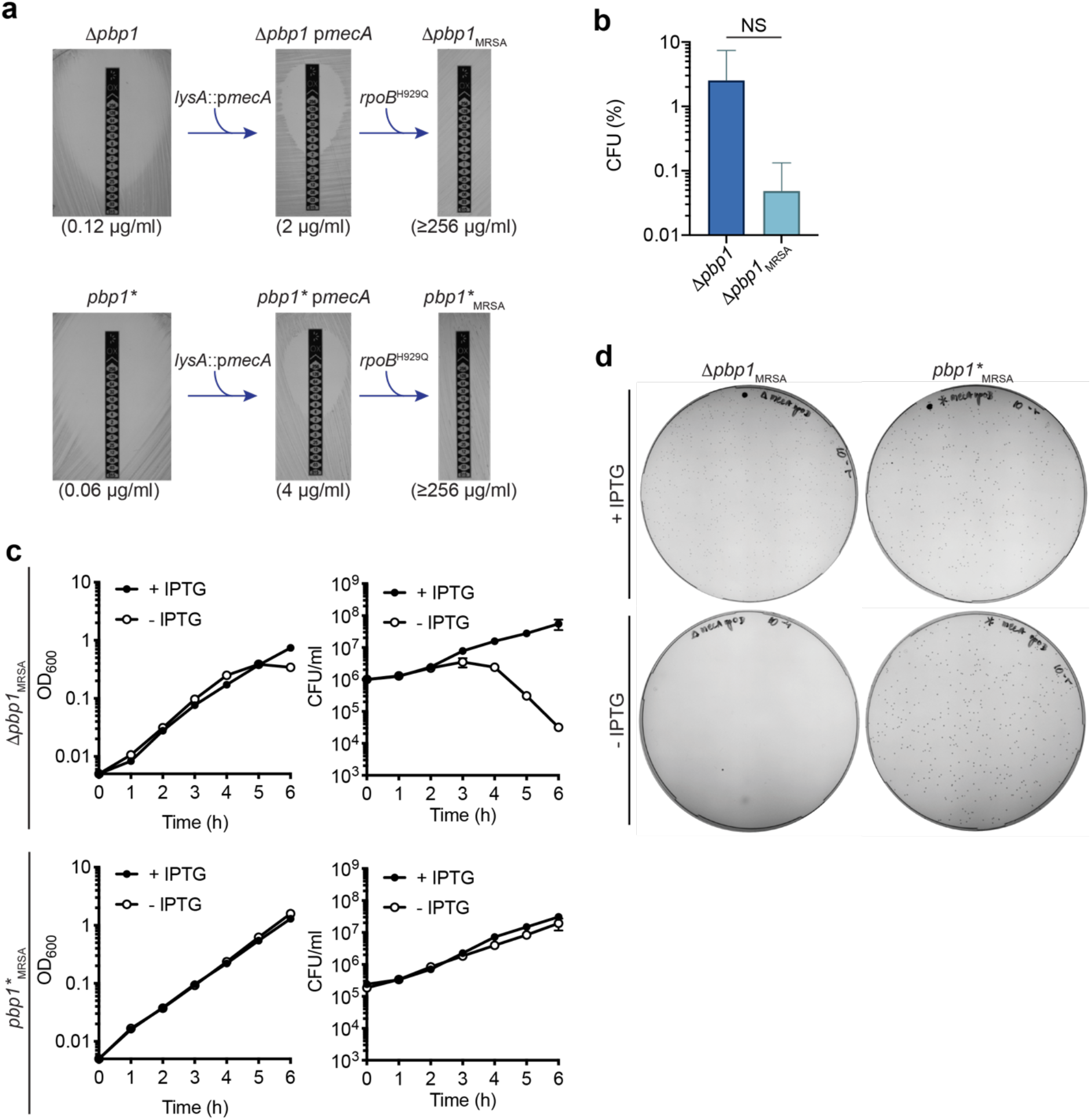
Essentiality of PBP1 and its TP activity in Methicillin-resistant *S. aureus*. **a**, Schematic representation of evolution of high-level *β*-lactam resistant Δ*pbp1* and *pbp1*.* A single copy *mecA* under its native promoter (p*mecA*) was introduced at the *lysA* locus, resulting in low-level oxacillin resistant Δ*pbp1* p*mecA* and *pbp1\** p*mecA.* Subsequently, addition of a point mutation in the *rpoB* gene results in development of high-level resistant Δ*pbp1*_MRSA_ and *pbp1\**. Oxacillin MICs shown in brackets were measured using the E-test strips. **b**, Plating efficiency of Δ*pbp* (MSSA) and MRSA Δ*pbp1*_MRSA_ cells upon the inducer removal compared to the control groups grown in the presence of inducer. Data represent mean ± SD. *P* value was determined by Mann–Whitney *U* tests. *P* = 0.5429 (NS, not significant). **c**, Growth curves of Δ*pbp1*_MRSA_ and *pbp1\**_MRSA_ in the presence or absence of IPTG. Data represent mean ± SD. Error bars that are smaller than the data point symbols are not shown. **d**, Growth of Δ*pbp1*_MRSA_ and *pbp1\**_MRSA_ with or without IPTG. Quantifications for Δ*pbp1*_MRSA_ are shown in Fig. 1 – figure supplement 2c and for *pbp1\**_MRSA_ are shown in Fig. 1b. Data are representative of at least three independent experiments.

**Fig. 2 – figure supplement 1.**
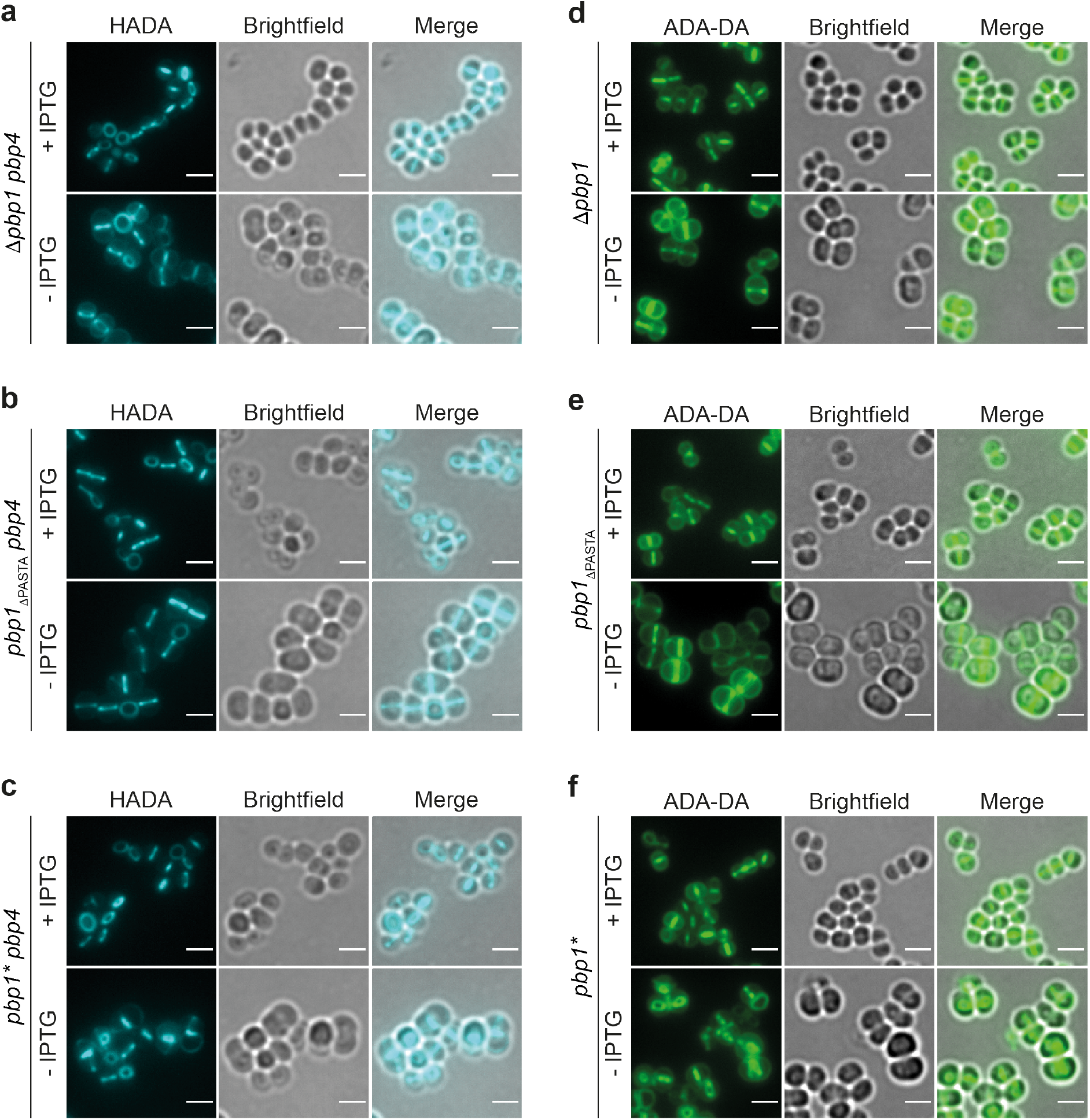
Loss of PBP1, PASTAs or TP activity of PBP1 does not prevent PG synthesis. **a-c**, PG incorporation in *pbp4* mutants depleted of PBP1. Δ*pbp1 pbp4, pbp1*_ΔPASTA_ *pbp4* and *pbp1* pbp4* grown with or without IPTG for 2 h and incubated with HADA for 5 min to show nascent PG incorporation. **d-f**, Δ*pbp1, pbp1*_ΔPASTA_ and *pbp1\** grown with or without IPTG for 2 h, incubated with dipeptide (ADA-DA) for 5 min and clicked to Alexa Fluor 488 to show nascent PG incorporation. Fluorescence images are average intensity projections of *z* stacks. Scale bars 2 μm. Images are representatives of two independent experiments.

**Fig. 2 – figure supplement 2.**
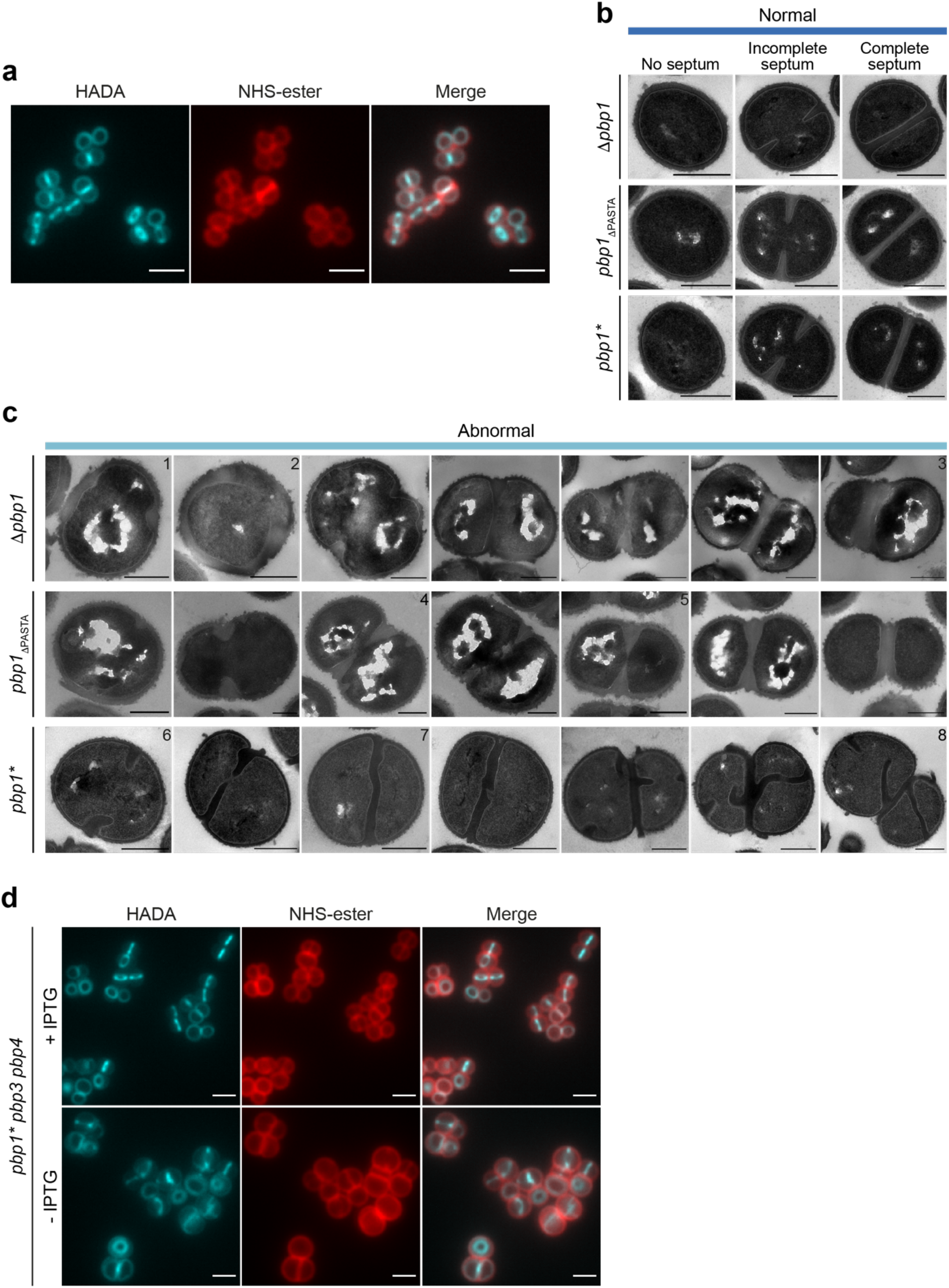
Role of PBP1 in *S. aureus*. **a**, Fluorescence images of SH1000 WT labelled with HADA for 5 min (nascent PG) and counter labelled with NHS-ester Alexa Fluor 555 (cell wall). Images are average intensity projections of *z* stacks. Scale bars 2 μm. **b**, TEM of Δ*pbp1, pbp1*_ΔPASTA_ and *pbp1\** grown in the presence of IPTG categorised as normal phenotype (blue). Scale bars 500 nm. **c**, TEM of Δ*pbp1, pbp1*_ΔPASTA_ and *pbp1\** grown in the absence of IPTG for 2 h categorised as abnormal phenotype (1, PG blebs; 2, thickened cell wall; 3, thickened complete septum; 4, multiple septa; 5, misshapen incomplete septum; 6, thick incomplete septum with rounded leading edge; 7, curved septum; 8, separation defect. Scale bars 500 nm. **d**, Fluorescence images of *pbp1* pbp3 pbp4* grown with or without IPTG for 2 h, labelled with HADA for 5 min (nascent PG) and counter stained with NHS-ester Alexa Fluor 555 (cell wall). Images are average intensity projections of *z* stacks. Scale bars 2 μm. Data are representative of two independent experiments.

**Fig. 2 – figure supplement 3.**
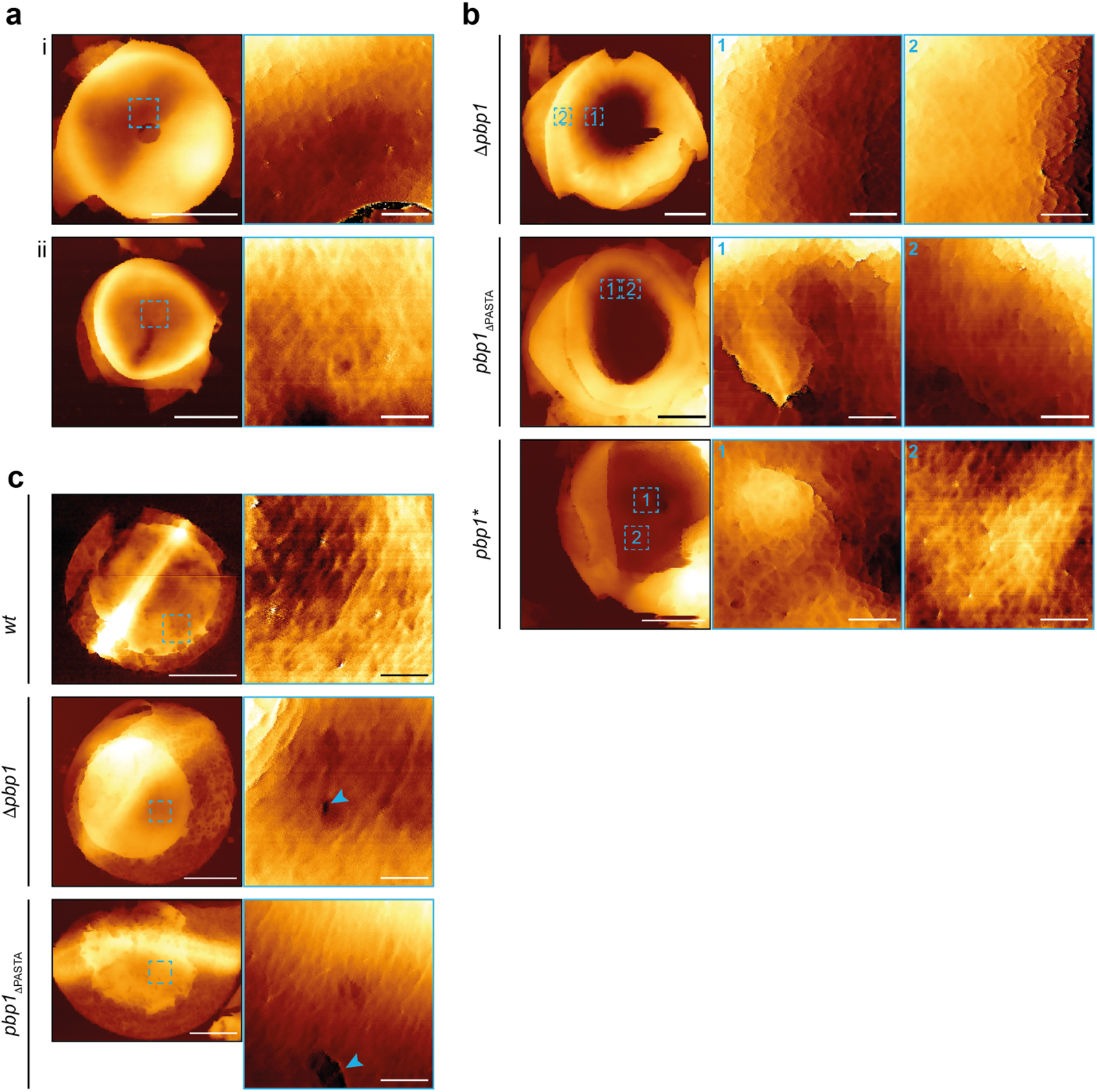
Gallery of AFM images of *S. aureus* Δ*pbp1, pbp1*_ΔPASTA_ and pbp1*. **a**, AFM topographic images of unfinished (i) and closed (ii) septa in *S. aureus* SH1000. Sacculi (images to the left, scale bars 500 nm, data scales (*z*): 200 (top) and 250 nm (bottom)) and higher magnification scans (images to the right, scale bars 50 nm, data scales (*z*): 80 (top) and 40 nm (bottom)) on the boxed areas from the images to the left. **b**, AFM topographic images of unfinished septa in Δ*pbp1* (from left to right: scale bars 500, 50 and 50 nm; data scales (*z*) 500, 120 and 150 nm)*, pbp1*_ΔPASTA_ (from left to right: scale bars 500, 50 and 50 nm; data scales (*z*) 693, 80 and 100 nm) and *pbp1\** (from left to right: scale bars 500, 50 and 50 nm; data scales (*z*) 500, 80 and 25 nm) grown in the absence of inducer for 2 h. Images to the left are sacculi, while images in the centre (1) and to the right (2) are higher magnification scans on the boxed areas of the images on the left. **c**, AFM topographic images of external nascent ring architecture in SH1000 WT (*wt*; scale bars: from left to right: scale bars 500 and 50 nm; data scales (*z*) 100 and 20 nm) and mutants Δ*pbp1* (scale bars: from left to right: scale bars 500 and 50 nm; data scales (*z*) 400 and 60 nm) and *pbp1*_ΔPASTA_ (scale bars: from left to right: scale bars 500 and 50 nm; data scales (*z*) 350 and 100 nm) grown in the absence of inducer for 2 h. Images to the left are sacculi, while images to the right are higher magnification scans on the boxed areas of the images on the left. The arrowheads indicate abnormal features, holes. Data are representative of two independent experiments.

**Fig. 3 – figure supplement 1.**
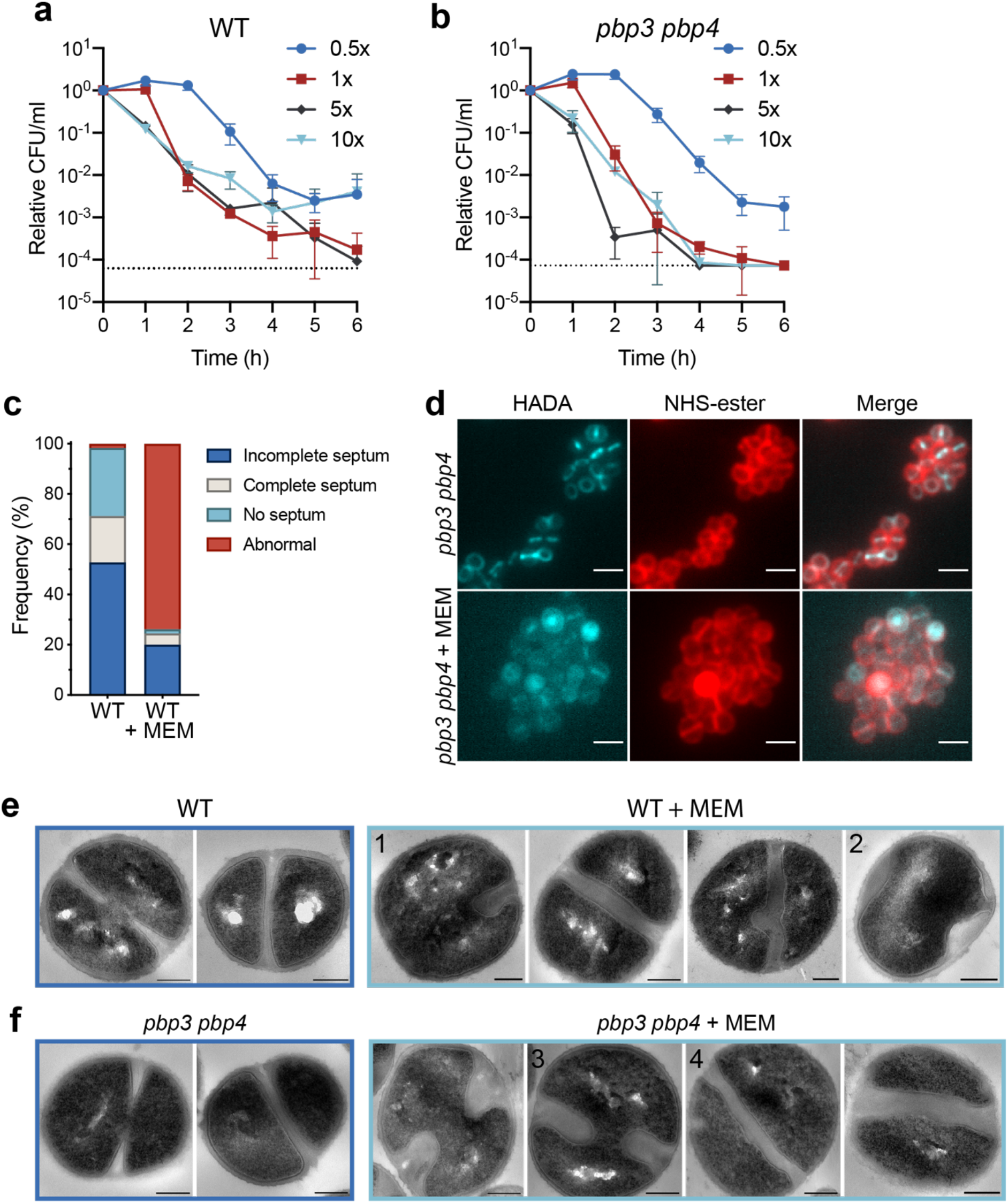
Effect of meropenem (MEM) on *S. aureus*. **a-b**, Bactericidal effect of addition of 0.5x, 1x, 5x and 10x MIC MEM on (a) SH1000 WT and (b) *pbp3 pbp4* (b). MEM MIC is 0.4 μg/ml and 0.2 μg/ml for SH1000 WT and *pbp3 pbp4*, respectively Data represent mean ± SD. Error bars that are symbols than the dots are not shown. The dotted line is the detection limit. **c**, Quantification of cellular phenotypes of SH1000 WT treated with 1x MIC MEM for 1 h based on HADA incorporation (Fig 3a). Same phenotype classification was used as shown in Fig. 2c. From left to right *n* = 309 and 355. **d**, Fluorescence images of *pbp3 pbp4* treated with 1x MIC MEM for 1 h, labelled with HADA for 5 min to show nascent PG and counter labelled with NHS-ester Alexa Fluor 555 (cell wall). Images are average intensity projections of *z* stacks. Scale bars 2 μm. Phenotype classification of MEM treated *pbp3 pbp4* was not possible due to low HADA fluorescence signal. **e-f,** TEM of SH1000 WT (e) and *pbp3 pbp4* (f) grown with or without 1x MIC MEM for 1 h. Scale bars 200 nm. Examples of cells categorised as normal phenotype are in blue, cells with abnormal phenotype are in light blue (1, asymmetric septum ingrowth; 2, off-septal PG thickening; 3, septum with rounded leading edge; 4, curved septum). Data are representative of three (a and b) and two (c) independent experiments. Experiments in d, e and f were performed once.

**Fig. 4 – figure supplement 1.**
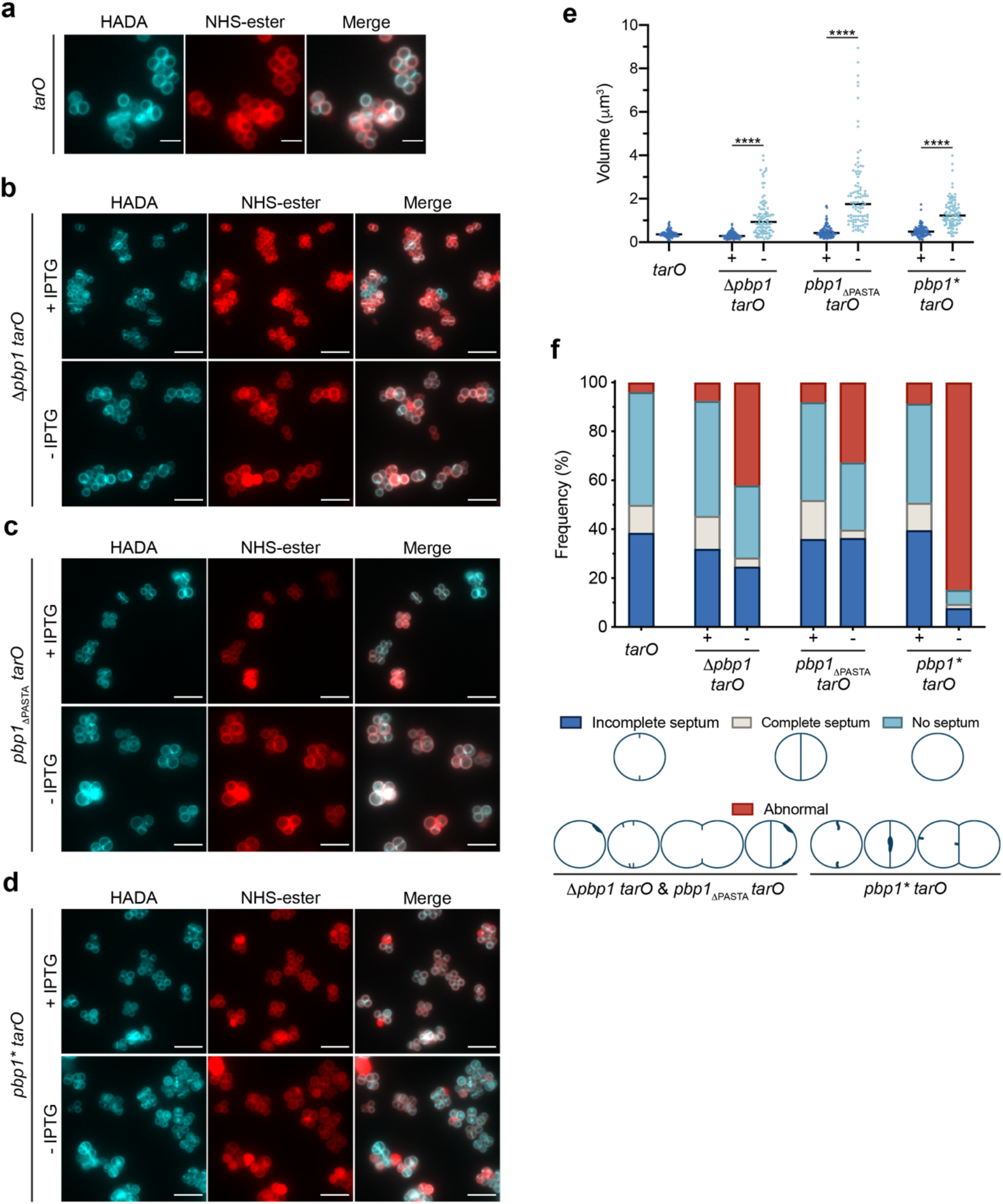
Functional association between PBP1 and WTA. **a**, Fluorescence images of the *tarO* mutant labelled with HADA for 5 min (nascent PG) and counter stained with NHS-ester Alexa Fluor 555 (cell wall). Images are average intensity projections of *z* stacks. Scale bars 5 μm. **b-d**, Δ*pbp1 tarO, pbp1*_ΔPASTA_ *tarO* and *pbp1* tarO* grown with or without IPTG for 2 h, incubated with HADA for 5 min to show nascent PG and counter labelled with NHS-ester Alexa Fluor 555 (cell wall). Images are average intensity projections of *z* stacks. Scale bars 5 μm. **e**, Cell volumes of *tarO* and Δ*pbp1 tarO, pbp1*_ΔPASTA_ *tarO* and *pbp1* tarO* grown with or without IPTG as measured by fluorescence microscopy after NHS-ester Alexa Fluor 555 labelling. Each dot represents a single cell. The median of each distribution is indicated by a black line. The number of cells analysed for each mutant and condition was *n* ≥ 100. *P* value was determined by Mann–Whitney *U* tests (***, *P* < 0.0001). From left to right: *P* = 2.243e- 022, 1.460e-037 and 8.074e-029; *n* = 100, 102, 101, 100, 100, 100 and 100. **f**, Quantification of cellular phenotypes for *tarO* and Δ*pbp1 tarO, pbp1*_ΔPASTA_ *tarO* and *pbp1* tarO* based on HADA incorporation (Fig. 4 – figure supplement 1a-d) after incubation with or without IPTG. From left to right *n* = 306, 253, 271, 358, 266, 313 and 336. Data are representative of two independent experiments.

**Fig. 5 – figure supplement 1.**
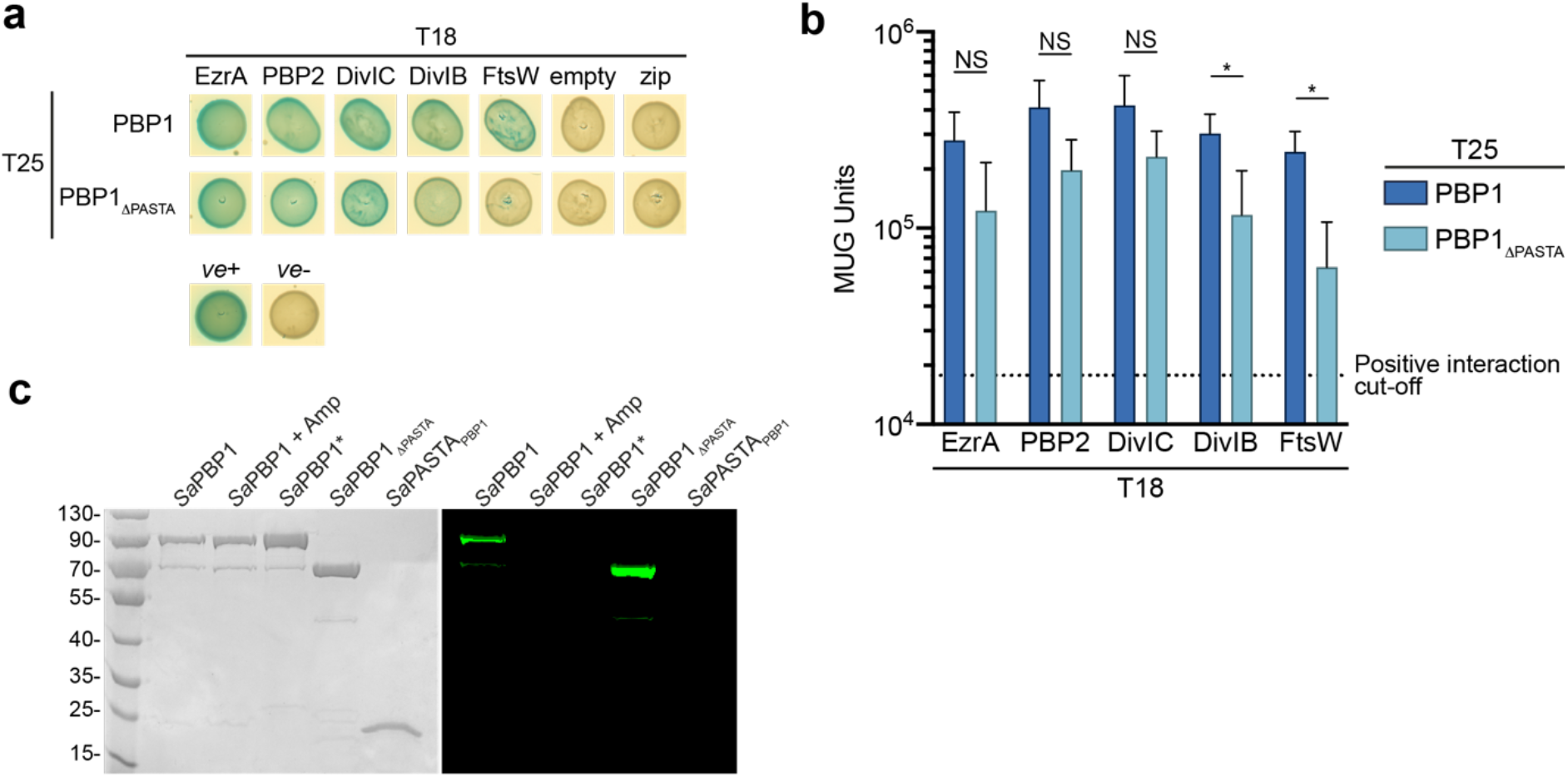
The role of PBP1 PASTA domains in interactions with cell division components and PG. **a**, Bacterial two-hybrid analysis of the effect of PASTA domains truncation on PBP1 interaction with its known interaction partners; empty, T18 with no insert; zip, T18 with a leucine zipper fragment; *ve*+, positive control (T18-zip/T25-zip); *ve*-, negative control (T18/T25). **b**, Quantitative bacterial two-hybrid analysis of the effect of the PASTA domains truncation on PBP1 interaction with cell division components determined by analysis of the β- galactosidase activities of *E. coli* BTH101 cells harbouring the corresponding plasmids. Dotted line, the positive interaction cut off value (4-fold greater than the pair of T18/T25). Data represent mean ± SD. *P* value was determined by Mann–Whitney *U* tests (*, *P* < 0.05). DivIB (PBP1 *vs* PBP1_ΔPASTA_) *P* = 0.0424, FtsW (PBP1 *vs* PBP1_ΔPASTA_) *P* = 0.0163. **c**, Coomassie-stained SDS-PAGE gel (left) and BocillinFL gel-based analysis (right) of purified recombinant *Sa*PBP1, *Sa*BPP1*, *Sa*PBP1_ΔPASTA_ and *Sa*PASTA_PBP1_. Bands corresponding to *Sa*PBP1 and *Sa*PBP1_ΔPASTA_ were fluorescent, indicating their covalent binding to BocillinFL. Bands corresponding to *Sa*PBP1* and *Sa*PASTA_PBP1_ were not fluorescent, and were therefore unable to bind BocillinFL. *Sa*PBP1 incubated with ampicillin prior to BocillinFL incubation failed to fluoresce, consistent with specific binding of BocillinFL to the TP domain. Expected sizes: *Sa*PBP1 and *Sa*PBP1*, 80.5kDa; *Sa*PBP1_ΔPASTA_, 64.5 kDa; *Sa*PASTA_PBP1_, 18.2 kDa. Data are representative of two (c) and three (a-b) independent experiments.

**Fig. 5 – figure supplement 2.**
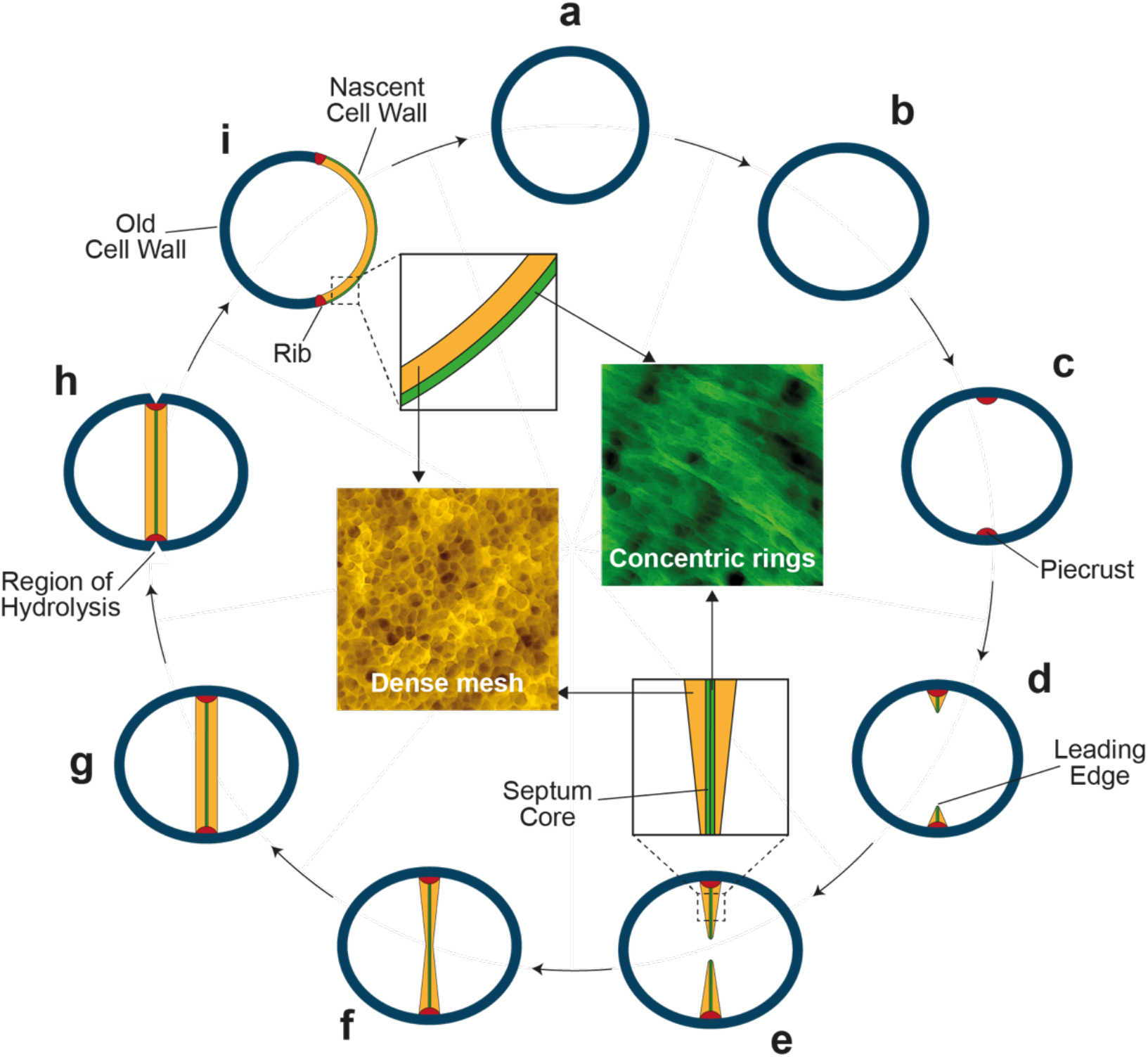
Conceptual model of septum formation in *S. aureus*. (a, b) The growing *S. aureus* cell increases in volume (Zhou et al., 2015). (c) Septal synthesis starts by formation of the piecrust (red) (Turner et al., 2010). (d. e) ‘V’ shaped septum (Lund et al., 2018) progresses inwards by insertion of ring like structured PG synthesised by PBP1- FtsW at the septum core and mesh structured PG produced by PBP2. (f) The annulus closes resulting in a bowed septum. (g) Septum is filled out by peptidoglycan insertion executed by PBP2 and this continues until the cross wall is of uniform thickness (Lund et al., 2018). (h) Cell wall is hydrolysed at the plane of septation. (i) Daughter cells separate. The cell wall of the daughter cell (coloured insets) is a chimera of the old cell wall with both internally and externally mesh structured PG and a nascent cell wall with the external ring structured PG and the mesh-like cytoplasmic facing PG (Pasquina-Lemonche et al., 2020).

## Appendix Tables

**Appendix Table 1.**
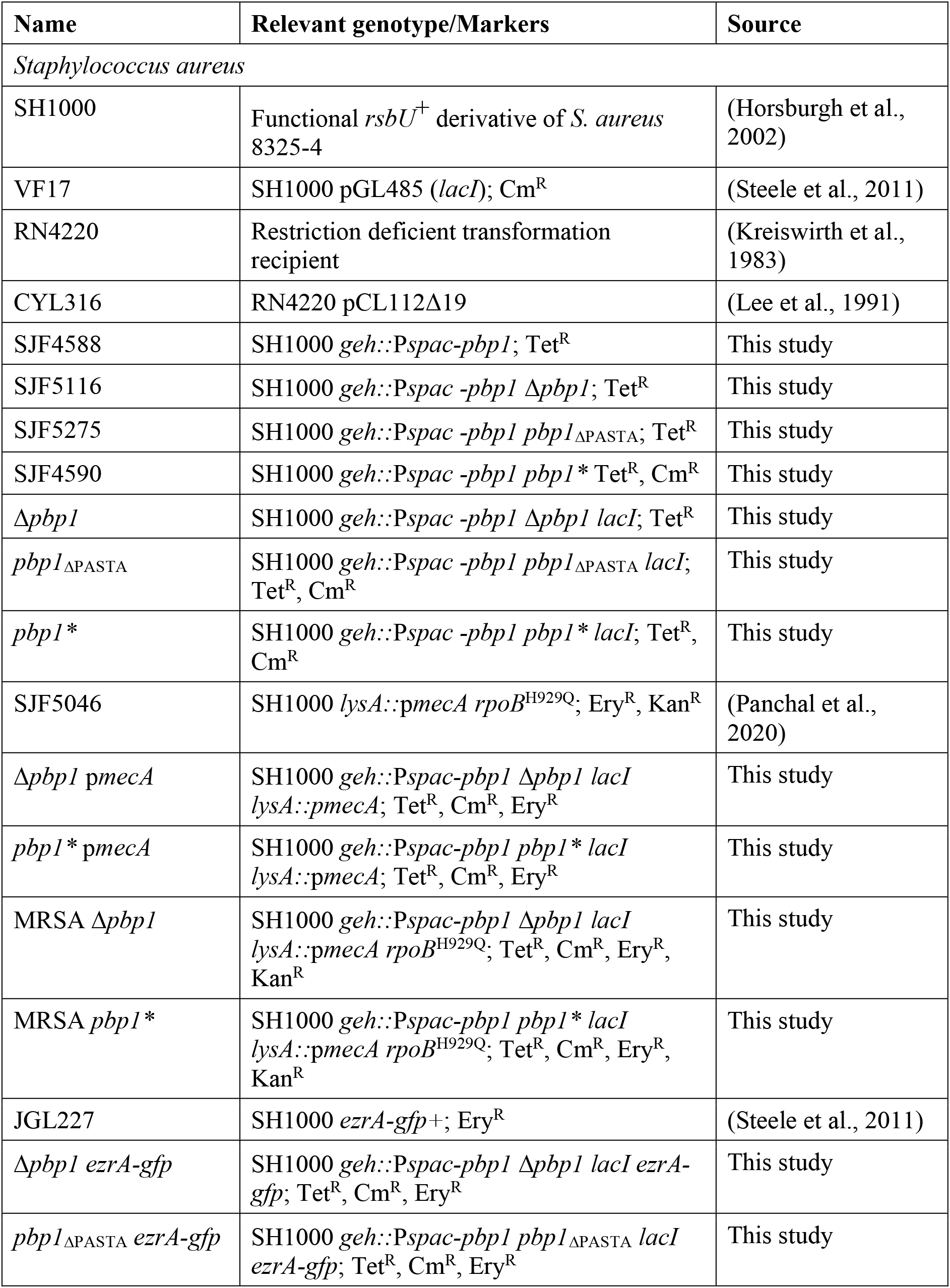
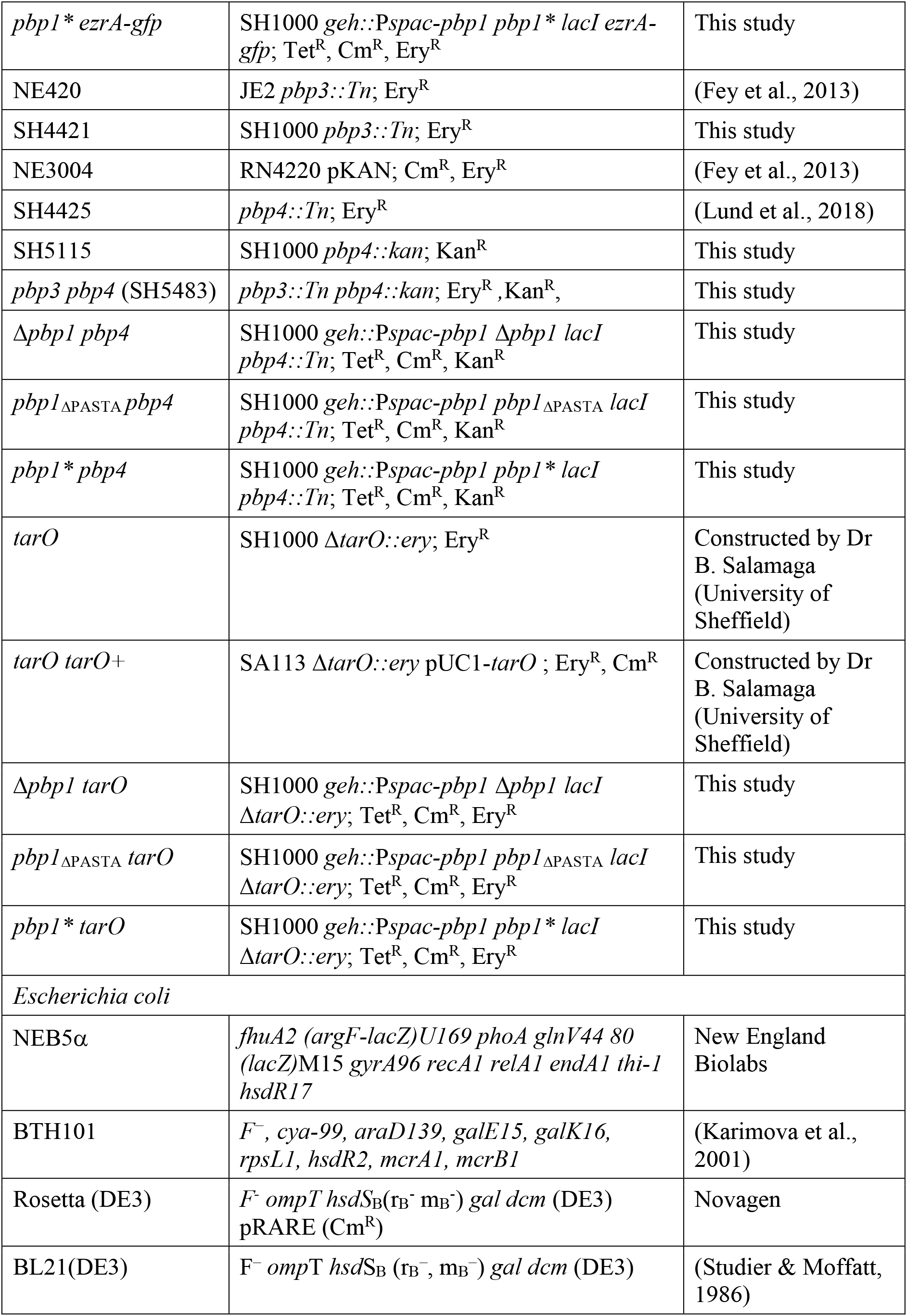
Strains used in this study.

**Appendix Table 2.** Plasmids used in this study.

| Name | Characteristics | Source |
| --- | --- | --- |
| pCQ11-FtsZ-SNAP | pCQ11 derivative containing *ftsZ-snap* under *Pspac*; Amp<sup>R</sup>, Ery<sup>R</sup> | (Lund et al., 2018) |
| pKASBAR | pUC18 containing *attP* and tetracycline cassette; Amp<sup>R</sup>, Tet<sup>R</sup> | (Bottomley et al., 2014) |
| pKB-*Pspac-pbpI* | pKASBAR containing *S. aureus pbpI* under *Pspac*; Amp<sup>R</sup>, Tet<sup>R</sup> | This study |
| pMAD | *E. coli-S. aureus* shuttle vector with temperature-sensitive origin of replication in *S. aureus* and constitutively produced thermostable β-galactosidase encoded by *bgaB*; Amp<sup>R</sup>, Ery<sup>R</sup> | (Arnaud et al., 2004) |
| pMAD-Δ*pbpI* | pMAD containing a deletion cassette for *S. aureus pbpI*; Amp<sup>R</sup>, Ery<sup>R</sup> | This study |
| pMAD-*pbpI*<sub>ΔPASTA</sub> | pMAD containing a deletion cassette for *S. aureus pbpI* PASTA domains; Amp<sup>R</sup>, Ery<sup>R</sup> | This study |
| pMAD-*pbpI*<sup>*</sup> | pMAD containing a cassette for introduction of a point mutation (S314A) in the active site *S. aureus pbpI*; Amp<sup>R</sup>, Ery<sup>R</sup> | This study |
| pGL485 | *E. coli-S. aureus* shuttle vector carrying *E. coli lacI* gene under the control of a constitutive promoter; Spec<sup>R</sup>, Cam<sup>R</sup> | (Cooper et al., 2009) |
| T18 (pUT18C) | Derivative of high copy-number pUC19, carrying gene encoding amino acids 225 to 399 of CyaA (T18 fragment); Amp<sup>R</sup> | (Karimova et al., 2001) |
| T18-zip (pUT18C-zip) | pUT18C coding for the leucine zipper region of the GCN4 yeast protein. Positive control; Amp<sup>R</sup> | (Karimova et al., 2001) |
| EzrA-T18 (pVF32) | pUT18(Karimova et al., 2001) containing T18 fused in frame to the 3' end of *S. aureus ezrA*; Amp<sup>R</sup> | (Steele et al., 2011) |
| T18-PBP2 (pGL547) | pUT18C containing T18 fused in frame to the 5' end of *S. aureus pbp2*; Amp<sup>R</sup> | (Steele et al., 2011) |
| T18-DivIC (pGL564) | pUT18C containing T18 fused in frame to the 5' end of *S. aureus divIC*; Amp<sup>R</sup> | (Steele et al., 2011) |
| T18-DivIB (pGL544) | pUT18C containing T18 fused in frame to the 5' end of *S. aureus divIB*; Amp<sup>R</sup> | (Steele et al., 2011) |
| T18-FtsW (pALB6) | pUT18C containing T18 fused in frame to the 5' end of *S. aureus ftsW*; Amp<sup>R</sup> | (Steele et al., 2011) |
| T25 (pKT25) | Derivative of low copy-number pSU40, carrying the first 224 amino acids of *B. subtilis* CyaA (T25 fragment); Kan<sup>R</sup> | (Karimova et al., 2001) |
| T25-zip (pKT25-zip) | pKT25 coding for the leucine zipper region of the GCN4 yeast protein. Positive control; Kan<sup>R</sup> | (Karimova et al., 2001) |
| T25-PBP1 (pGL550) | pKT25 containing T25 fused in frame to the 5' end of *S. aureus pbp1*; Kan<sup>R</sup> | (Steele et al., 2011) |
| T25-PBP1<sub>ΔPASTA</sub> | pKT25 containing T25 fused in frame to the 5' end of *S. aureus pbp1*<sub>ΔPASTA</sub> (M1-S595); Kan<sup>R</sup> | This study |
| pOPINRSF | *kan P<sub>T7</sub> lacI*; Kan<sup>R</sup> | (Berrow et al., 2009) |
| pVR01 | *kan P<sub>T7</sub> pbp1 lacI*; pOPINRSF derivative for overexpression of full length *S. aureus* PBP1 (1-744); Kan<sup>R</sup> | This study |
| pVR02 | *kan P<sub>T7</sub> pbp1(37-744) lacI*; pOPINRSF derivative for overexpression of *Sa*PBP1 (37-744); Kan<sup>R</sup> | This study. |
| pVR03 | *kan P<sub>T7</sub> pbp1\*(37-744: S314A) lacI*; pOPINRSF derivative for overexpression of *Sa*PBP1* (37-744); Kan<sup>R</sup> | This study |
| pVR04 | *kan P<sub>T7</sub> pbp1(37-595) lacI*; pOPINRSF derivative for overexpression of *Sa*PBP1<sub>ΔPASTA</sub>(37-595); Kan<sup>R</sup> | This study |
| pVR06 | *kan P<sub>T7</sub> pbp1(595-744) lacI*; pOPINRSF derivative for overexpression of *Sa*PASTA<sub>PBP1</sub> (595-744); Kan<sup>R</sup> | This study |
| pOPINJB | *bla P<sub>T7</sub> lacI*; Amp<sup>R</sup> | (Berrow et al., 2009) |
| pSA50 | pOPINJB derivative for overexpression of sPBP1A-BAP; Amp<sup>R</sup> | This study |

**Appendix Table 3.**
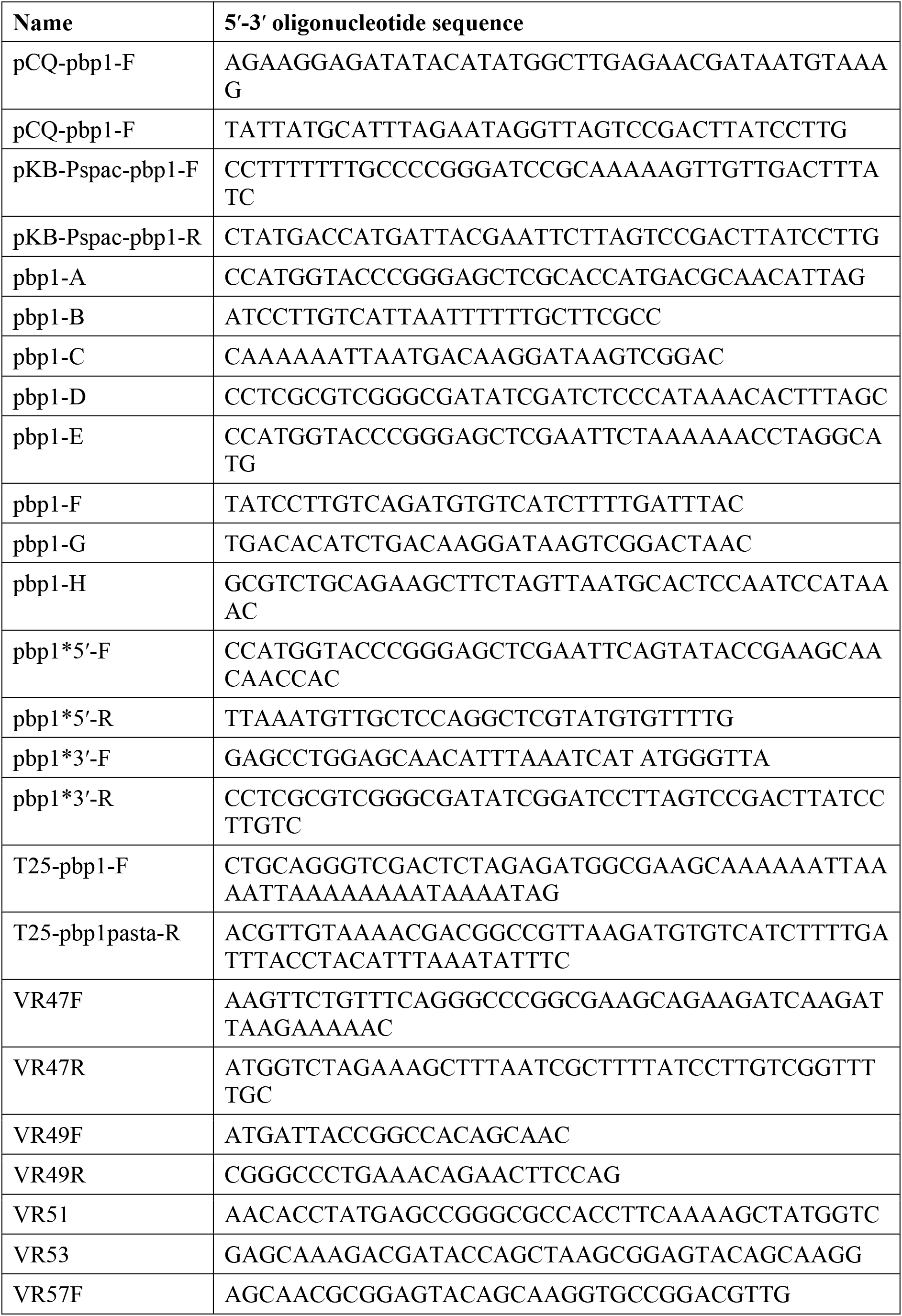
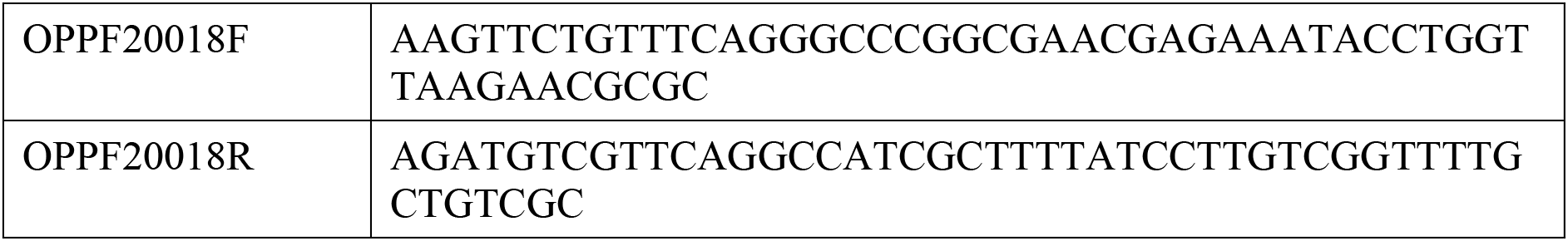
Oligonucleotides used in this study.

**Appendix Table 4.**
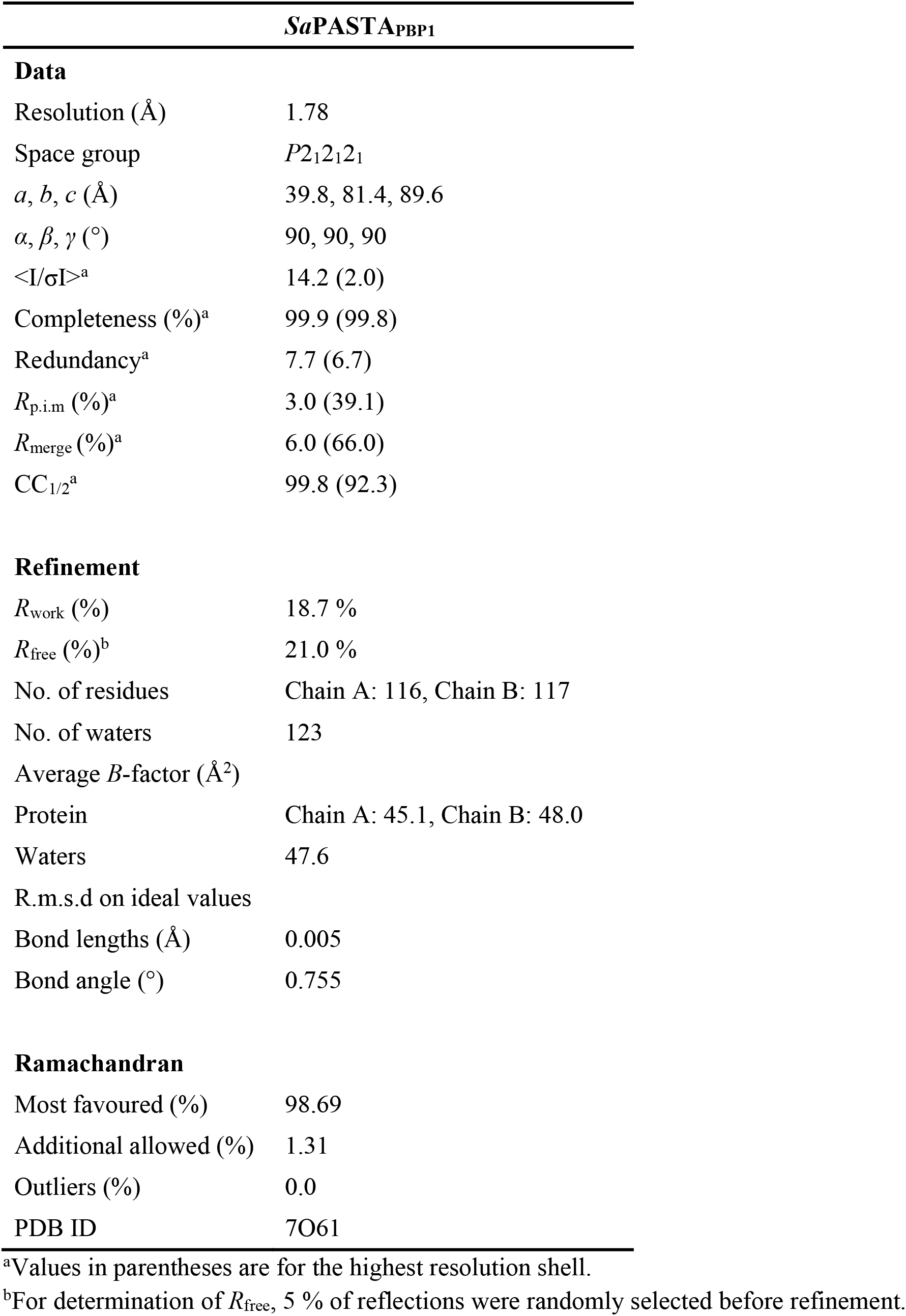
Crystallographic data.

